# Generation of hypoimmunogenic gastric insulin-secreting organoids

**DOI:** 10.64898/2026.07.08.736836

**Authors:** Anna A. Dattoli, Matthew E. Brown, Bradley Pearson, Zachary Feinstein, Ying Lan, Vasumati Polavarapu, Max Zhou, Ryan Nachman, Yosip Kelemen, Shahin Rafii, Remi J. Creusot, Todd M. Brusko, Qiao Zhou, Xiaofeng Huang

## Abstract

Gastric insulin-secreting organoids (GINS) represent a promising source of β-like cells for type 1 diabetes (T1D) therapy. In same-donor comparisons with induced pluripotent stem cell-derived islets (iPSC-islets), GINS displayed robust glucose responsiveness and reduced expression of key T1D autoantigens. Importantly, GINS exhibited decreased susceptibility to cytotoxicity mediated by engineered HLA-matched preproinsulin-specific effector T cells (Avatar Teffs) and a distinct transcriptional profile enriched for immune-modulatory and stress-adaptive gene programs. To enhance immune evasion, we engineered gastric stem cells to overexpress Programmed Death Ligand 1 (PD-L1) in an inducible manner. PD-L1^+^ GINS maintained normal functionality, while exhibiting improved survival under allogeneic Avatar Teff challenge in a MHC class I-independent fashion. We evaluated PD-L1-mediated protection against autologous Avatar Teff attack using an endothelialized microfluidic platform recapitulating physiologic immune interactions. T cells show reduced infiltration into PD-L1⁺ GINS, resulting in significantly higher organoid viability compared to control GINS. Together, these findings identify GINS as a functional and engineerable β-like cell platform with intrinsic hypoimmunogenic features, and support PD-L1 engineering as a strategy to enhance immune protection for both allogeneic and autologous transplantation in T1D.

## Introduction

Type 1 diabetes (T1D) results from autoimmune destruction of insulin-producing β-cells, leaving patients dependent on lifelong exogenous insulin. While insulin therapy has transformed diabetes from a fatal disease into a manageable condition^1^, insulin injections remain an imperfect solution, often leading to glycemic instability, hypoglycemic episodes, and long-term complications^2^. Emerging β-cell replacement therapies, whether from cadaveric donors (e.g., Lantidra, FDA.gov) or stem cell-derived sources (e.g., Vx-880, Clinical Trial NCT04786262), offer promising avenues for restoring glucose homeostasis and preventing life-threatening hypoglycemic episodes^3^. However, their clinical utility is limited by the requirement for lifelong immunosuppression, which poses serious risks and excludes many patients from eligibility^4,5^.

To address these challenges, autologous transplantation strategies using direct cellular reprogramming offer a compelling alternative, naturally circumventing immune rejection (reviewed by Dattoli et al.^6^). Specifically, our group has developed a novel approach using autologous gastric insulin-secreting organoids (GINS), reprogrammed from the patient’s own gastric stem cells, that can be generated in just 10 days and have reversed diabetes in mouse models^7^. GINS represent a unique, autologous β-cell source that avoids alloimmune rejection. However, autoimmune attack remains a major obstacle in T1D. For instance, although GINS can survive for over six months in immunocompromised mice^7^, their survival in immune-competent hosts with active autoimmunity remains unproven.

Multiple research efforts are exploring immune evasion strategies to create β-cells that are resistant to T cell-mediated autoimmune and innate immune killing. One such strategy involves engineering β-cells to express CD47, a surface protein that binds to SIRPα on macrophages and NK cells, delivering a “don’t-eat-me” signal that prevents phagocytosis and cytotoxicity^8^. This approach has advanced to early clinical evaluation, including trials testing CD47-engineered human islets^9^. Another approach focuses on incorporating immune-inhibitory molecules to counteract both cellular and humoral immune responses. In this context, the Deuse lab showed that synthetic engagers with agonistic functionality to their inhibitory receptors, such as TIM-3 and SIRPα, can effectively protect engineered HLA-deficient stem cell-derived endothelial cells from NK cell and macrophage-mediated killing^10^. Furthermore, inclusion of a truncated version of CD64 helps shield the grafted cells from antibody-dependent cellular cytotoxicity^9^. Interestingly, Programmed Death-Ligand 1 (PD-L1, CD274) is a surface molecule broadly expressed on somatic cells and plays a central role in maintaining immune tolerance^11^. By engaging its receptor, Programmed Death-1 (PD-1), which is expressed on antigen-stimulated T cells, PD-L1 dampens inflammatory responses and limits CD8⁺ T cell cytotoxicity. This inhibitory interaction prevents excessive T cell activation against self-antigens and is critical for immune homeostasis. Tumor cells frequently exploit this pathway as an adaptive mechanism to evade anti-tumour immunity^12^. In the context of islet transplantation, enhancing PD-1/PD-L1 signalling within the local microenvironment represents a promising strategy to protect β-cells from immune-mediated destruction. Indeed, recent studies have shown that induction of PD-L1 overexpression, alone or in combination with HLA class I knockout, protects stem cell-derived β-cells from diabetogenic CD8^+^ T cell attack^13,14^. However, this approach might still require suppressing innate immune responses against HLA-deficient cells^8,10^.

In the present study, our scRNA-seq data show that GINS expressed significantly lower levels of major T1D autoantigens, including glutamic acid decarboxylase 65 (GAD65)^15^ and zinc transporter 8 (ZnT8)^16^, compared to primary human islets. This suggests that they might be naturally less prone to autoimmunity. To validate this hypothesis, we carried out differentiation of induced pluripotent stem cells (iPSCs) and gastric stem cells (GSCs) obtained from the same donor into islet-like organoids (iPSC-islets and GINS, respectively). While GINS and iPSC-islets exhibited comparable insulin content, GINS showed a reduced polyhormonal cell fraction and robust glucose responsiveness. In addition, single-cell RNA-seq analyses revealed lower expression of key T1D autoantigens in GINS. In same-donor comparisons, GINS also displayed reduced susceptibility to cytotoxicity mediated by preproinsulin (PPI)-specific effector T cells (Avatar Teffs), consistent with intrinsic hypoimmunogenic features. To enhance immune protection, we engineered GSCs via inducible cell surface overexpression of PD-L1. Importantly, this genetic modification did not impact functionality of PD-L1^+^ GINS compared to unmodified GINS (Ctrl GINS). Strikingly, PD-L1 overexpression reduced cell death upon incubation with either allogeneic or autologous Avatar Teffs. Importantly, to model human T1D disease in a dynamic setting, we further developed an endothelialized microfluidic platform that enables visualization and quantification of a T cell-mediated response against GINS in 3D. Collectively, our data identify GINS as a high-functioning with robust glucose-stimulated insulin secretion, intrinsically hypoimmunogenic, and readily engineerable β-cell platform, with enhanced immune protection through PD-L1 overexpression. These findings support further development of GINS for immune-protected cell replacement strategies in T1D.

## Results

### GINS demonstrate improved functional performance compared with iPSC-islets obtained from the same donor

Our previous study demonstrated that GSCs can give rise to insulin-secreting, glucose-responsive GINS with high efficiency compared to human islets^7^. Furthermore, scRNA-seq revealed that GINS organoids lack GAD2 (GAD65) and show low expression of SLC30A8 (ZnT8), two major T1D autoantigens^17^ (Fig. S1A). We next sought to verify whether these properties were maintained when evaluated alongside stem cell-derived β-cells obtained from the same donor. To enable a unique matched same-donor human comparison while minimizing inter-donor variability, we obtained stomach and spleen tissue from a single deceased organ donor. Access to paired stomach and lymphoid tissues from the same organ donor is rare, making this matched design a valuable platform for controlled benchmarking across cell sources. Therefore, GSCs were isolated from the stomach corpus as previously shown^7^ and genome-edited to express inducible sequential expression of Ngn3 followed by Mafa/Pdx1 (as detailed in the methods). In parallel, splenocytes were reprogrammed into iPSCs (Fig. 1A). iPSCs were then tested for pluripotency markers OCT3/4, NANOG and SOX2 (Fig. S1B). Using previously established methods^7,18,19^, we performed directed differentiation of iPSCs and GSCs into iPSC-islets and GINS (Fig. 1A). We next performed immunofluorescence (Fig. 1B) and flow cytometry analyses to assess expression of the β-cell marker C-peptide (C-PEP). While no differences in the proportion of C-PEP⁺ cells were observed at Day 19 for GINS or Day 34 for iPSC-islets (Fig. 1B and 1C), follow-up analysis within approximately one week revealed a divergence between the two systems: GINS exhibited increased C-PEP expression, whereas iPSC-islets showed a decline in expression of the same marker (Fig. 1C). Moreover, our flow confirms the presence of mature MAFA⁺C-PEP⁺ cells in GINS organoids compared with iPSC-islets (Fig. 1D). In addition, the fraction of polyhormonal cells expressing both C-PEP and glucagon (GCG) was drastically reduced in GINS (34-fold), indicating a greater purity of insulin-secreting cells derived from GSCs (Fig. 1E and S1C). To gain deeper insights into the gene expression profiles of the two samples during β-like cell differentiation, we performed scRNA-seq on mature iPSC-derived islets at day 34 and GINS at day 19 of their respective differentiation protocols. Our combined heat map shows the distribution of iPSC-islets and GINS population (Fig. 1F). We first identified the INS expressing region within the two types of organoids (Fig. 1F). We then resolved multiple cell populations within the organoids, including β-like, α-like and δ-like endocrine cells, as well as smaller non-endocrine populations, including epithelial-like, endothelial-like, fibroblast-like and proliferative cells (Fig. 1G). Within the β-like compartment, sub clustering identified *INS*-high and *INS*-low populations. Notably, *INS*-high cells were showing reduced glucagon- and somatostatin-positive cells and diminished representation of off-target non-endocrine identities, while being enriched for markers associated with β-cell maturity, including INS, IAPP, MAFA (which is overexpressed in GINS), UCN3 and G6PC2 (Fig. 1G and S1D-E). These observations suggested that *INS*-high cells represent a more mature β-like state present in both GINS and iPSC-islets. We then conducted a more stringent analysis, comparing the expression of other β-cell identity markers between GINS and iPSC-islets in both whole organoids and the *INS*-high fraction. This analysis revealed increased expression of multiple β-cell identity and functional markers in GINS, including INS, GCK, ABCC8, PAX6, NKX2-2, PCSK1 and G6PC2^7^, both at the whole-organoid level and within the INS-high compartment, supporting enrichment of a more mature β-like transcriptional program in GINS (Fig. 1H). To assess functional properties of GINS and iPSC-islets, we performed static glucose-stimulated insulin secretion (GSIS) assays. While iPSC-islets secreted higher absolute insulin levels under these conditions (Fig. 1I), GINS exhibited a robust glucose-responsive secretory profile, as reflected by an increased stimulation index. Together, these findings indicate that both platforms support insulin production, while suggesting that GINS harbor features consistent with a more mature glucose-responsive β-like state.

**Fig. 1.**
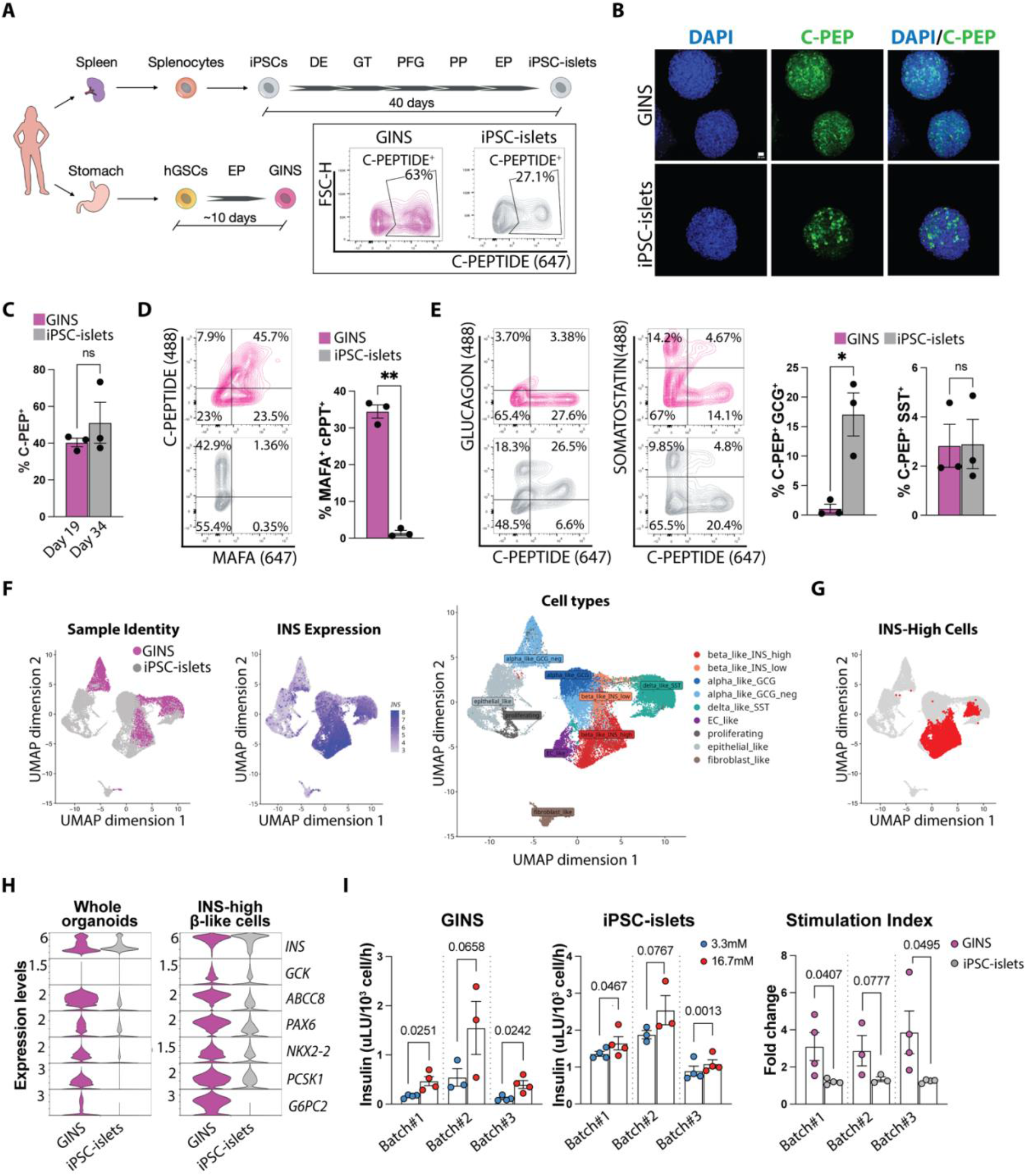
GINS show better glucose responsiveness compared to iPSC-islets upon differentiation. (A) Schematic of β-like cell differentiation using iPSC-derived islets and/or GINS from the same donor and flow cytometry plots highlighting C-PEPTIDE^+^ fraction. (B) Immunofluorescence staining for C-peptide (C-PEP, green) and DAPI (blue) at day 18 for GINS and day 34 for iPSC-islets. Scale bars represent 20 μm. (C) Flow cytometry quantification of C-PEP expression on days 19 and 28 for GINS and days 34 and 41 for iPSC-islets. (D) Flow cytometry plots and quantification of C-PEP and MAFA expression on day 18 for GINS and day 34 for iPSC-islets. (E) Flow cytometry quantification of Glucagon (GCG) and Somatostatin (SST) in combination with C-PEP expression on day 28 for GINS and day 41 for iPSC-islets. (F) UMAP of GINS organoids and or iPSC-islets showing INS expression and different cell types identified. (G) UMAP of GINS organoids and or iPSC-islets showing INS high fraction. (H) Violin plots from scRNA-seq of selected genes in β-like cells obtained either from GINS or iPSC-islets in both whole organoids and in the INS high fraction. (I) GSIS of GINS and iPSC-islets at day 16 and day 42 of their respective protocol. All graphs plot mean and Standard Error of the Mean (SEM). All quantifications are based on n=3 independent differentiations (unless differently specified) and were analyzed by paired t test.

### GINS exhibit a hypoimmunogenic phenotype with reduced vulnerability to T cell attack

Violin plots from scRNA-seq analysis revealed reduced expression of key T1D autoantigens, including *GAD2* (GAD65) and *SLC30A8* (ZnT8), in GINS compared to human islets (Fig. S1D). We next asked whether a similar pattern could be observed when comparing GINS to iPSC-islets. To this end, we generated a T1D autoantigen module score based on established targets, including *GAD2*, *SLC30A8*, *PTPRN* (IA-2), *ICA1*, *G6PC2* (IGRP), and *IAPP* (Fig. 2A). This analysis did not reveal a significant difference between GINS and iPSC-islets at the aggregate level (Fig. 2A). However, gene-level analysis (Fig. 2B) revealed a more nuanced pattern: while expression of several islet-associated antigens, such as *PTPRN* and *ICA1*, was comparable or higher in GINS, that of *GAD2* and *SLC30A8*, two well-established and clinically relevant T1D autoantigens, was selectively reduced. Together, these findings suggest that GINS exhibit a distinct antigenic profile characterized by reduced expression of specific dominant T1D autoantigens. To validate these findings, we performed flow cytometry analysis assessing GAD65 and ZnT8 expression within the C-PEP⁺ population.

**Fig. 2.**
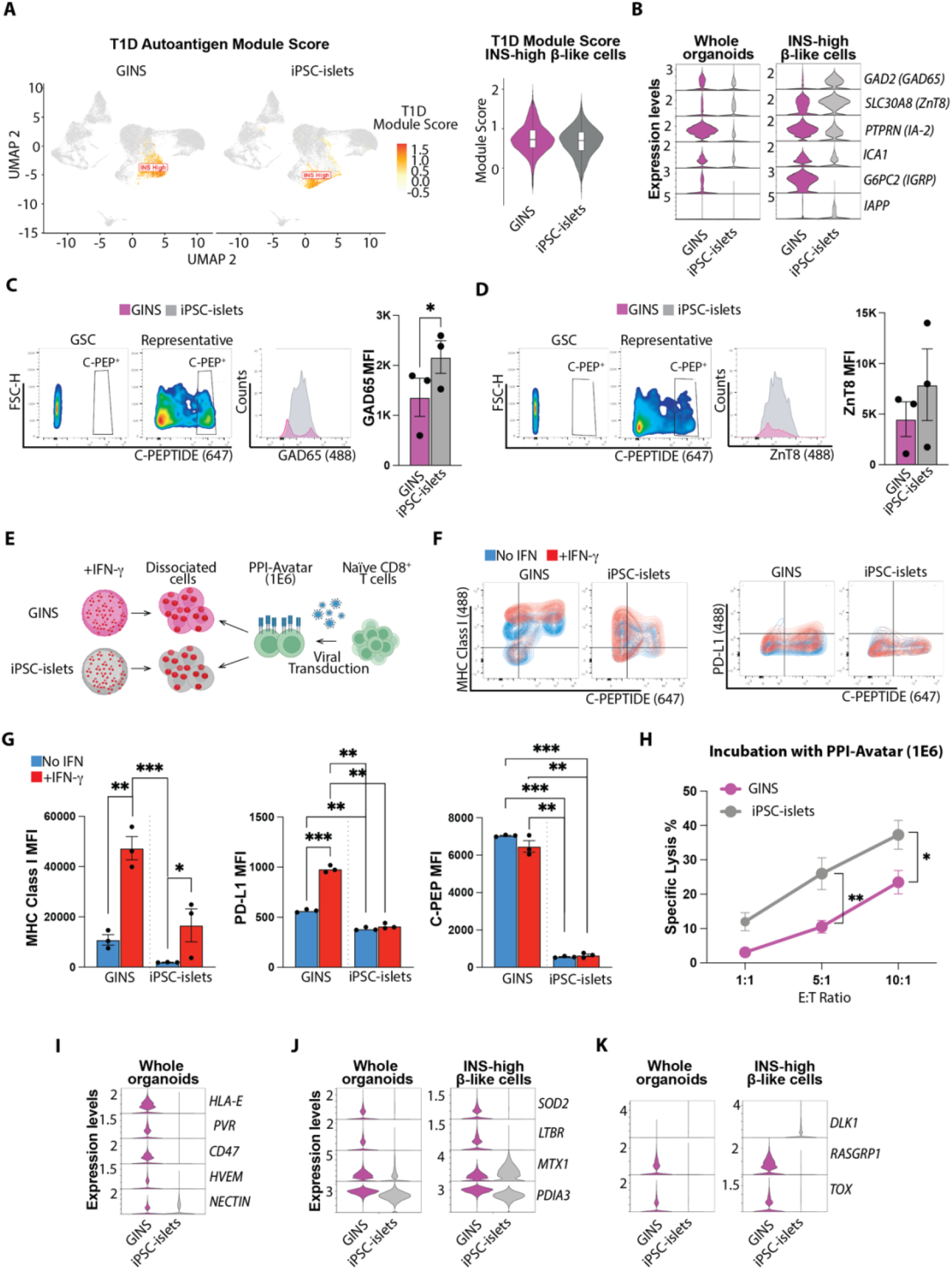
GINS are naturally hypoimmunogenic compared to iPSC-islets. (A) UMAP of T1D module autoantigen score and violin plots of the total T1D module score obtained either from GINS or iPSC-islets in the INS high fraction. (B) Violin plots from scRNA-seq of classic β-cells autoantigens obtained either from GINS or iPSC-islets in both whole organoids and in the INS high fraction. (C) Flow cytometry plots of C-PEP, GAD65, ZnT8 expression on day 19 for GINS and day 34 for iPSC-islets. (D) Flow cytometry quantification of C-PEP, GAD65, ZnT8 expression on day 19 for GINS and day 34 for iPSC-islets. (E) Schematic for allogeneic immune assay with GINS and iPSC-islets. (F) Flow cytometry quantification of C-PEP, MHC class I, PD-L1 at day 23 for GINS and day 41 for iPSC-islets upon 24 hrs IFN-γ treatment. (G) Flow cytometry plots of MHC class I and or PD-L1 expression in combination with C-PEP. (H) Cell death quantification measured as % specific lysis of GINS (Day 23) and iPSC-islets (Day 43) upon incubation with allogeneic T cells specific for preproinsulin peptide (PPI-Avatar 1E6). (I) Violin plots from scRNA-seq of classic immune evasion genes obtained either from GINS or iPSC-islets in whole organoids. (J) Violin plots from scRNA-seq of inflammatory stress-related genes of T1D pancreas and pancreatic lymph nodes^30^ obtained either from GINS or iPSC-islets in both whole organoids and the INS high fraction. (K) Violin plots from scRNA-seq of β-cell death-related genes^35^ obtained either from GINS or iPSC-islets in both whole organoids and the INS high fraction. All graphs depict mean and SEM. All quantifications are based on n=3 independent differentiations (unless differently specified) and were analyzed by paired t test. *p < 0.05; **p < 0.01; ***p < 0.001.

Consistent with the scRNA-seq results, flow cytometry demonstrated a 1.6-fold reduction in GAD65 protein levels within the C-PEP⁺ high population of GINS relative to iPSC-islets. A similar trend was observed for ZnT8, albeit not significant (Fig. 2C-D). Since our results revealed reduced expression of islet autoantigens in GINS compared to iPSC-islets (and human islets), we reasoned that GINS may be inherently less susceptible to autoimmune attack. Moreover, as GINS are gastric insulin-secreting cells, their molecular distinctions compared to iPSC-islets could further mitigate immunogenicity. To test this hypothesis, we assessed the survival of GINS and iPSC-islets following immune challenge with cytotoxic CD8^+^ T cells (Teffs) specific for the preproinsulin (PPI) β-cell antigen (PPI-Avatar) (Fig. 2E, schematics). During T1D pathogenesis, IFN-γ plays a central role in promoting immunogenicity by upregulating MHC-I molecules, thereby enhancing antigen presentation to Teffs^20,21^. To model this environment, we pre-incubated GINS and iPSC-islets with varying doses of IFN-γ and observed that 6 ng/mL for 24 hrs was sufficient to induce robust MHC-I expression in both GINS and iPSC-islets, while higher concentrations did not further augment expression, as shown by flow cytometry (Fig. 2F-G and Fig. S1F). Since IFN-γ can also induce PD-L1 expression and thereby promote immune regulation^22^, we assessed PD-L1 protein levels in GINS and iPSC-islets before and after treatment (Fig. 2F-G). At baseline, GINS exhibited ∼1.3-fold higher PD-L1 expression compared to iPSC-islets. Following IFN-γ exposure, we observed no significant change in the proportion of PD-L1-expressing cells within iPSC-islets (Fig. 2F-G). In contrast, GINS displayed a twofold elevation in PD-L1 expression, as measured by mean fluorescence intensity (MFI), upon IFN-γ treatment (Fig. 2G). We also observed an increased fraction of PD-L1⁺ C-PEP⁺ cells in GINS following IFN-γ treatment (Fig. S1G), an effect that was not observed in iPSC-islets. Importantly, overall C-PEP expression remained unchanged (Fig. 2G), indicating that the increased PD-L1⁺ C-PEP⁺ fraction was driven by upregulation of PD-L1 rather than changes in β-cell identity or abundance. We then isolated HLA-A2-matched naïve Teffs and transduced them with a T cell receptor (TCR) reactive to the PPI15-24 peptide^23^, a canonical T1D autoantigen, creating PPI-reactive Avatar Teffs (PPI-avatar, clone 1E6). We then co-cultured dissociated iPSC-islets and GINS with PPI-Avatar. As negative control, we used Avatar Teffs specific for an irrelevant antigen (MART-1, clone DMF5^24^), and as a positive control, we used CAR Teffs specific for HLA-A2. Cells were incubated with each Avatar Teffs at the following Effector:Target (E:T) ratios: 1:1, 5:1, 10:1 (Fig. 2H and S2A). After 48 hrs, MART-1 Teffs induced minimal cell death, whereas HLA-A2 CAR Teffs drove robust lysis, validating the system (Fig. S2A). Notably, when co-cultured with PPI-specific Avatar Teffs, GINS exhibited significant reduced susceptibility to T cell-mediated killing compared to iPSC-islets. At an E:T ratio of 5:1, GINS showed an approximately 70% reduction in specific lysis, which remained evident at 10:1 with an approximately 47% reduction (Fig. 2H). Together, these findings suggest that GINS display intrinsic features consistent with reduced immunogenicity. To further explore this phenotype, we performed transcriptomic analyses (Fig. 2I-K). GINS showed higher expression of several genes associated with immune modulation, including HLA-E, PVR, CD47, HVEM, and NECTIN (Fig. 2I)^25–28^. Importantly, PVR (CD155) was already found out to protect β-cell from cytotoxic attack^29^. In parallel, analysis of gene sets informed by spatial transcriptomic studies of pancreatic tissue and draining lymph nodes in T1D^30^ indicated increased expression of genes associated with inflammatory-stress adaptation, including SOD2, LTBR, MTX1 (Fig. 2J)^31–34^. Finally, guided by recent work identifying DLK1/MEG3, RASGRP1, and TOX as β-cell-mediated T1D risk loci^35^, we found that GINS showed higher expression of RASGRP1 and TOX but lower expression of DLK1 compared with iPSC-islets (Fig. 2K), consistent with a transcriptional state associated with β-cell survival and maturation. Collectively, these findings support that GINS exhibit a distinct transcriptional profile combining immune-modulatory and stress-adaptive features, which may contribute to their reduced susceptibility to T cell-mediated cytotoxicity.

### PD-L1 overexpression preserves β-like cell identity and functionality

Although GINS cells exhibited greater immune evasion than iPSC-islets, they were not completely resistant to immune attack (Fig. 2H). Notably, our scRNA-seq analysis suggests that this partial resistance may be driven by a distinct molecular profile, including enrichment of immune evasion gene signatures (Fig. 2I). Based on this, we sought to determine whether the well-established coinhibitory receptor PD-L1^12^ could further enhance GINS protection against immune attack. Indeed, previous work has shown that PD-L1 can protect stem cell-derived β-cells in HLA-knockout contexts^13,14^. Therefore, we engineered GSCs with a Tet-On inducible system, enabling sequential induction of *Ngn3ER* under the 4OH-Tamoxifen treatment followed by a doxycycline (Dox)-inducible *MAFA-PDX1-PD-L1* construct under the TRE promoter, using our previously established lentiviral transduction protocol^7^ (Fig. 3A, simplified scheme). PD-L1-TetOn GSCs maintained expression of canonical GSC markers such as SOX9, while they are still dividing, as shown by the expression of Ki67, compared to unmodified controls (Fig. 3A). Upon differentiation into GINS, organoids maintain their morphology and immunofluorescence analysis showed comparable C-PEP signal in both control and PD-L1⁺ GINS, while PD-L1 expression was restricted to PD-L1⁺ GINS (Fig. 3B). Because our *Ngn3ER*-hGSCs were labelled with constitutive mCherry (see Materials and Methods for further details on the construct), PD-L1 expression was quantitatively assessed in engineered GINS by flow cytometry, gating on the mCherry reporter (PD-L1⁺ mCherry⁺) and confirmed by immunofluorescence (Fig. 3C-D). We further validated comparable expression of MAFA (Fig. 3E and S2B), as well as hormones including C-PEP, GCG, and somatostatin (SST) (Fig. S2C) in both organoid types. Additionally, control and PD-L1⁺ GINS displayed similar functional responses upon glucose stimulation (Fig. S2D and 3F), demonstrating that inducible PD-L1 overexpression does not impair β-like cell function. Finally, we also did not observe a significant difference in the expression levels of GAD65 and ZnT8 between the organoids (Fig. S2E-F-G). Taken together, these findings demonstrate that GINS represent an engineerable cell source and that PD-L1 overexpression preserves gastric-derived β-like cell morphology, identity, and functionality.

**Fig. 3.**
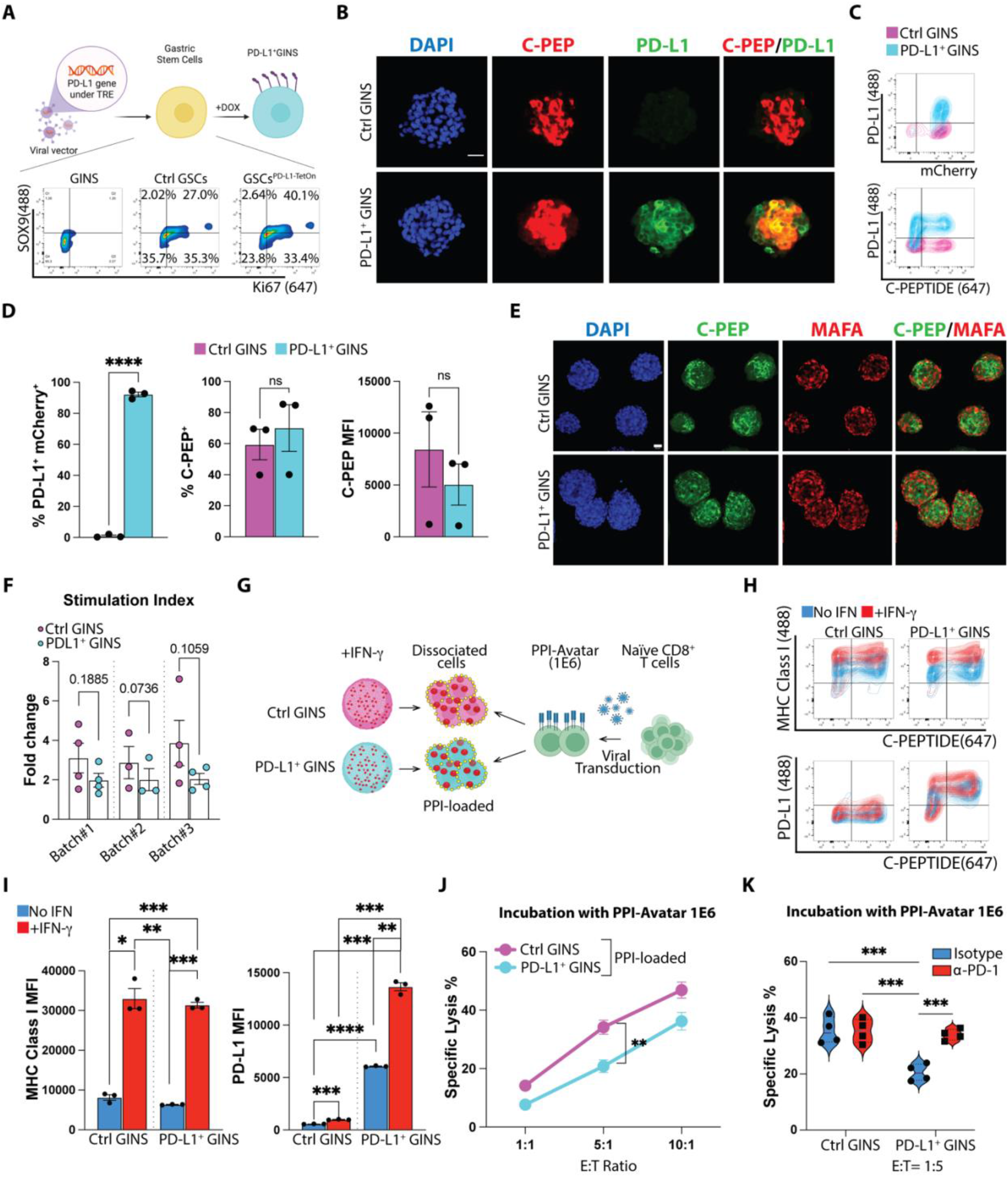
PD-L1 protect GINS against cytotoxic allogeneic T cell attack. (A) Schematic for lentivirus genome recombination for insertion of PD-L1 under TRE promoter into Ctrl GSC line and flow cytometry of GSC markers SOX9 and cell division marker Ki67. (B) Wide field images of immunofluorescence staining for DAPI (blue), C-PEP (red), PD-L1 (green), in Ctrl GINS and PD-L1^+^ GINS at day 16 upon differentiation at 20× magnification. Scale bars represent 50 μm. (C) Flow cytometry plots of PD-L1 expression in combination with mCherry reporter and C-PEP in Ctrl and PD-L1^+^ GINS at day 18. (D) Flow cytometry quantification of PD-L1, mCherry, C-PEP expression. (E) Confocal images of immunofluorescence staining for DAPI (blue), C-PEP (green), MAFA (red), in Ctrl GINS and PD-L1^+^ GINS at day 16 upon differentiation. Scale bars represent 20 μm. (F) Stimulation Index from GSIS assay comparing Ctrl GINS and PD-L1^+^ GINS at day 16. (G) Schematic for allogeneic immune assay with PPI-loaded Ctrl GINS and PD-L1^+^ GINS. (H) Flow cytometry plots of MHC class I and or PD-L1 in combination with C-PEP at day 23 for Ctrl GINS and PD-L1^+^ GINS upon 24 hrs IFN-γ treatment. (I) Flow cytometry quantification of MHC class I molecules and PD-L1 measured as MFI in Ctrl and PD-L1^+^ GINS at day 18. (J) Cell death quantification measured as % specific lysis of Ctrl GINS and PD-L1^+^ GINS (Day 23) upon incubation with allogeneic T cells specific for preproinsulin peptide. All graphs show mean and SEM. All data were analyzed by paired t test. ns, not significant. (K) Cell death quantification measured as % specific lysis of Ctrl GINS and PD-L1^+^ GINS (Day 23) upon coculture with allogeneic T cells specific for preproinsulin peptide previously incubated with either Isotype or PD-1 antibody. All graphs show mean and SEM. All quantifications are based on n=3 independent differentiations (unless differently specified) and by paired t test. ns, not significant; *p < 0.05; **p < 0.01; ***p < 0.001; ****p < 0.0001.

### PD-L1 overexpression enhances immune protection of GINS against antigen-specific allogeneic T cells

To determine whether PD-L1 confers additional immune resistance against cytotoxic CD8⁺ T cells (Fig. 3G, schematics), we employed the immune challenge assay described above (Fig. 2G). Prior to co-culture, we verified that IFN-γ stimulation (6 ng/mL) induced comparable upregulation of MHC-I molecules in both groups (Fig. 3H-I). As expected, IFN-γ treatment also led to ∼2.5-fold increase in PD-L1 expression in PD-L1^+^ GINS (Fig. 3I), without significantly altering the fraction of C-PEP⁺ PD-L1^+^ cells (Fig. S2H). We next performed a chromium release assay^36^ to evaluate cytotoxicity (Fig. 3J and S2I). Given our prior observation of baseline resistance of GINS to T cell-mediated killing, we hypothesized that this could reflect inefficient presentation of the PPI peptide. Such limited antigen presentation might obscure the protective effect of PD-L1 overexpression evaluated by this assay. To address this, we loaded both control and PD-L1^+^ GINS organoids with PPI ^37^. Organoids were then co-cultured with PPI-Avatar, MART-1-Avatar, or HLA-A2 CAR Teffs at varying E:T ratios for 48 hours (Fig. 3J and S2H). Cell-mediated lysis was used as a measure of cytotoxicity. As expected, MART-1-Avatar induced minimal cell death at all ratios, whereas HLA-A2 CAR Teffs induced the highest levels of cell death (Fig. S2H). We observed higher baseline levels of cell death in GINS in this assay compared to the previous experiment (Fig. 2G and 3J), consistent with the increased antigen-specific T cell challenge introduced by PPI15-24 peptide loading. Importantly, we observed a significant ∼37% reduction in cell lysis when comparing control GINS to PD-L1⁺ GINS incubated with PPI-specific Avatar Teffs at the 5:1 ratio (Fig. 3J). Reductions of ∼40% and ∼15% were also observed at the 1:1 and 10:1 ratios, respectively, though these were not statistically significant. Together, these findings indicate that PD-L1 expression enhances the immune-evasive properties of GINS. To confirm that the observed protection was specifically mediated by PD-L1 signalling rather than genetic background differences introduced by viral transduction, we examined the contribution of the PD-L1:PD-1 axis. PPI-Avatar T cells were pre-treated with an anti-PD-1 blocking antibody and cocultured with either control or PD-L1⁺ GINS (Fig. 3K) at the 1:5 E:T ratio. Compared with isotype control treatment, PD-1 blockade did not significantly alter T cell cytotoxicity toward control GINS. In contrast, blockade of PD-1 during coculture with PD-L1⁺ GINS resulted in significantly increased target cell death compared with the isotype control (Fig. 3K). These results demonstrate that the protective effect conferred by PD-L1 overexpression in GINS is functionally dependent on PD-L1:PD-1 signalling.

### PD-L1 overexpression mitigates cytotoxicity in a human Organ-on-Chip Model

Our previous results showed that PD-L1 overexpression is well tolerated by GINS (Fig. 3) and enhances their immune evasion from PPI-specific T cells derived from an allogeneic donor (Fig. 3). We next decided to further evaluate whether PD-L1 can protect GINS in a more complex system designed to model human T1D. Although mouse models have been instrumental in advancing our understanding of T1D^38,39^, transplantation of multiple organoids into a single graft site complicates quantitative assessment of early events such as vascularization^40^ and T cell infiltration. For this reason, we employed an Organ-on-Chip model perfused with autologous Avatar Teffs, enabling the first *in vitro* modelling of T1D on a microfluidic system (Fig. 4A). Using this system, we observed vascularization supported solely by endothelial cells, without other tissue/cell contribution, and quantified T cell infiltration at the single organoid level. For this purpose, naïve CD8⁺ T cells were isolated from the splenocytes, and gastric stem cells isolated of the same deceased donor were used to generate both control (Ctrl) GINS and PD-L1⁺ GINS (Fig. 4A). Splenocytes from our donor contained approximately 16% naïve CD8⁺ T cells (CD45RA^+^), gated on live singlets, which were then enriched to ∼98% purity (Fig. S3A). Similarly to our previous work with allogeneic Avatar-Teffs, naive CD8⁺ T cells were transduced with a lentiviral construct encoding the PPI-specific TCR linked via a T2A sequence to GFP (Fig. S3B). GFP⁺ cells were then sorted to generate PPI Teffs. Importantly, 55% of these cells expressed PD-1 upon stimulation with rhIL-2 and rhIL-7 (Fig. S3C). Control T cells (Ctrl Teffs) were instead transduced with GFP alone. We then perfused organoids with autologous PPI-Avatar-Teffs or Ctrl Teffs with an endothelialized microfluidic device that recapitulates *in vivo* vascularization (Fig. 4A-B-C). Specifically, the endothelial platform previously validated for supporting vascularization of human islets^41^, was here extended to model autoimmune interactions in a controlled *in vitro* environment. To assess the feasibility of this approach, we first used iPSC-islets, that recapitulate canonical β-cell identity, representing the established targets of autoreactive PPI Teffs, generated from the same donor splenocytes used to isolate autologous T cells. Following IFN-γ stimulation, 50 iPSC-islets organoids (approximately 2,000 cells/each) were stained with Vybrant Dil and co-cultured with Human Umbilical Vein Endothelial Cells (HUVECs). Cell mix was seeded into the device, achieving robust vascularization within 48 hrs (Fig. 4A and S3D-D’). PPI-Avatar-Teffs or Ctrl Teffs (200,000 T cells/sample) were then introduced and co-cultured for 48 hours (Fig. 4B). Imaging revealed that Ctrl Teffs remained dispersed across the device without specific target engagement (Fig. 4B, panels I-I’), whereas PPI-Avatar-Teffs localized intensely to the organoids, showing a pronounced degree of infiltration (Fig. 4B, panels II, II’). Cell death was quantified by flow cytometry (Fig. 4B and S3E-F). After dissociating the organoids and matrix, endothelial cells were stained for CD31, a classical endothelial marker^42^, Annexin V to label initial apoptotic cells^43^, and DAPI to label dead cells. Because both T cells and endothelial cells were GFP⁺, this strategy enabled us to gate on GFP⁻ cells, thereby selectively isolating the organoid population for analysis (Fig. S3E-E’-E’’). As shown in Fig. 4B and S3F, iPSC-islets incubated with PPI-Avatar-Teffs showed a ∼1.4-fold reduction in Annexin V⁻ DAPI⁻ cell frequency compared with those incubated with Ctrl Teffs, across two technical replicates. This suggests that iPSC-islets are susceptible to autologous T cell attack and validates our immune-challenge platform. We next extended this assay to Ctrl and PD-L1⁺ GINS (Fig. 4C-F). Because the assay spans five days, we first sought to confirm that GINS maintained insulin expression throughout the co-culture period. Accordingly, insulin expression was assessed after six days of co-culture with endothelial cells, revealing sustained C-PEPTIDE expression under these conditions (Fig. 4C). We then moved to our previously validate experimental set up. Therefore, following IFN-γ stimulation, using the same conditions as the previous experiment with iPSC-islets, organoids were seeded onto the endothelialized platform, which vascularized within 48 hours. (Fig. 4D, panels I-I’). We then perfused autologous PPI-Avatar-Teffs or Ctrl Teffs into each sample (Fig. 4D panels II to III’). Consistent with observations in iPSC-islets, Ctrl Teffs remained dispersed throughout the device (Fig. 4D, panels II-II’). Notably, PPI-Avatar-Teffs infiltrated Ctrl GINS (Fig. 4D, panel III), but infiltration of PD-L1⁺ GINS was reduced (Fig. 4D, panel III’), which we quantified using an infiltration index (Fig. 4E). Specifically, GFP fluorescence intensity was measured inside and outside each organoid, and infiltration was expressed as the ratio of inside to outside organoid GFP signal (Fig. S3H). As shown in Fig. 4E, PPI-Avatar-Teffs infiltrated control GINS approximately threefold more than Ctrl Teffs. In contrast, PPI-Avatar-Teffs showed a comparable infiltration index to Ctrl Teffs with PD-L1⁺ GINS. Additionally, flow cytometry confirmed the PD-L1 protective effect, revealing ∼1.2-fold higher survival of PD-L1⁺ GINS relative to Ctrl GINS when challenged with PPI Teffs (Fig. 4F). Together, these findings indicate that, within our endothelialized microfluidic platform, PD-L1 overexpression attenuates infiltration and confers immune protection to GINS against autologous cytotoxic T cells.

**Fig. 4.**
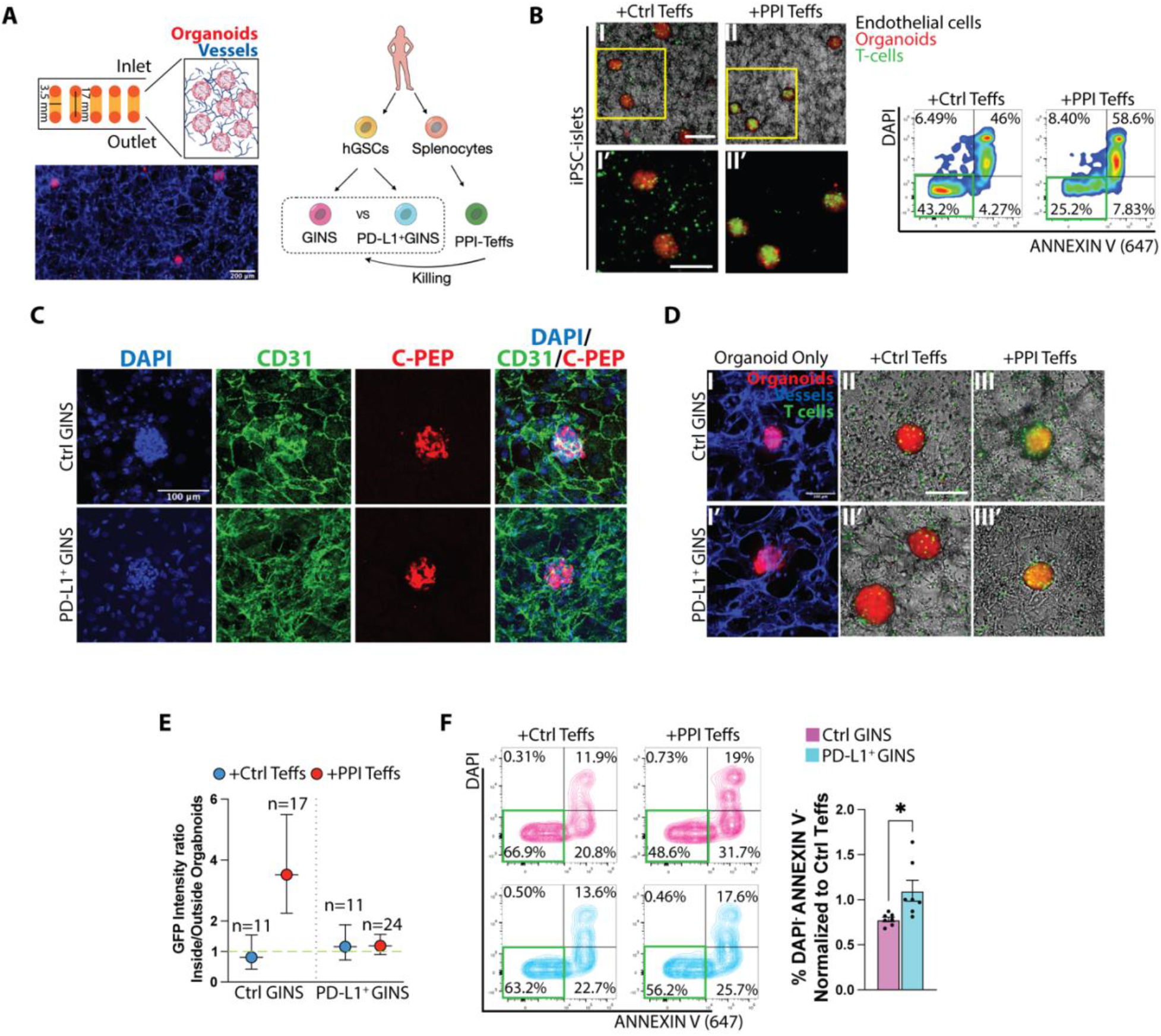
PD-L1 protects GINS against cytotoxic autologous T cell attack. (A) Schematics of the endothelial microfluidic device used to vascularize organoids and schematic of autologous immune challenge with Ctrl GINS and PD-L1^+^ GINS, scale bar represents 200 μm. (B) Wide field images of iPSC-islets alone or incubated with autologous Ctrl Teffs (GFP^+^ only), and PPI-Teffs (also GFP^+^) on the endothelial platform and flow cytometry of cell viability upon T cell incubation measured using DAPI and Annexin V, scale bar represents 100 μm. (C) Immunofluorescence staining for CD31 (green), C-peptide (C-PEP, red) and DAPI (blue) at day 20 for Ctrl GINS and PD-L1^+^ GINS. Scale bars represent 100 μm. (D) Panels I-I’: Confocal images of vascularized organoids; Panels II-II’-III-III’: Wide field images of Ctrl GINS and PD-L1^+^ GINS incubated with autologous Ctrl Teffs (GFP^+^ only) and PPI-Teffs on the endothelial platform, scale bar represents 100 μm. (E) Infiltration Index quantification; n = number of organoids analysed. Error bars represent the confidence interval at 95%. (F) Flow cytometry and relative quantification of cell viability measured using DAPI and Annexin V (*n* = 3 biological replicates, 7 technical replicates). All graphs depict mean and SEM, unless otherwise specified. All quantifications are based on n=3 independent differentiations (unless otherwise specified) and were analyzed by paired t test. ns, not significant; *p < 0.05; **p < 0.01; ***p < 0.001; ****p < 0.0001.

## Discussion

The success of a long-lasting pancreatic β-cell replacement therapy is tightly linked to immune tolerance to mitigate immune rejection and autoimmune destruction. Therefore, developing immune protection of the graft is crucial. Previous approaches, based on HLA knockout, such as β2-microglobulin (for HLA class I) or CIITA (for HLA class II), can effectively block antigen presentation of the transplanted stem cell-derived β-cell^14,27,44^, protecting against T cell cytotoxicity, but HLA null cells are susceptible to innate immune rejection. Targeted depletion of HLA class I and class II alongside CD47 overexpression protects human islet grafts against innate and T cell-mediated killing^9^. These previous works show that the concept of immunological protection can effectively inhibit immune response and guarantee graft survival. However, these approaches require HLA class I and II knockout.

Here, we show that compared to same-donor iPSC-derived islets, GINS show robust functionality and a lower burden of most prevalent and clinically relevant T1D autoantigens. Importantly, our results indicate that GINS are intrinsically (about 70%) less susceptible to killing by PPI-specific effector T cells (Fig. 1-2). Future directions should investigate the mechanism of T cell inhibition in terms of T cell cytokine release, as well as immune evasion mechanisms employed by GINS. Importantly, GINS are not β-cells, and our data suggest that they may harbor a unique intrinsic immune-evasion signature, including transcriptional factors and surface molecules that dampen antigen presentation and effector engagement. Additionally, GINS might be more resistant to cytokine stress (Fig. 2). Systematic mapping of this signature (by multi-omics/CRISPR perturbation) could reveal transferable targets such as antigen-processing nodes and/or adhesion/co-stimulation modulators. These nodes could be transferred to strengthen stem cell-derived β-cells and/or protect human islets against immune attack.

To further enhance immune evasion, we engineered GSCs to overexpress PD-L1 under an inducible system. PD-L1⁺ GINS preserved glucose-responsive function relative to unmodified controls yet showed superior survival against both allogeneic and autologous Avatar-Teff challenge (Fig. 3-4), consistent with reports that PD-L1 shields stem cell-derived β-cells from T cell attack^13,14^. Notably, in our system PD-L1 overexpression confers an additional ∼20-30% reduction in T cell-mediated cytotoxicity across assays. This modest yet consistent effect may reflect the intrinsically reduced susceptibility of GINS to immune attack, thereby limiting the magnitude of additional protection that can be achieved through PD-L1 alone. Nonetheless, these data indicate that PD-L1 functions as an effector-phase checkpoint that synergizes with GINS’ intrinsically low immunogenicity. Different from previous findings, in our allogeneic setting, this strategy mitigates the need for HLA knock-out approaches, offering a simpler, potentially safer manufacturing route. Notably, at higher E:T ratios (10:1), PD-L1⁺ GINS remain susceptible to T cell-mediated cytotoxicity, indicating that immune evasion is partial and does not result in complete escape from immune surveillance.

We leveraged a HUVEC endothelialized microfluidic perfusion platform that recreates a perfusable capillary bed, recapitulating immune-β-like organoid interactions under physiologically relevant conditions. Our endothelialized platform allows the longitudinal monitoring of the vascularization of organoids by the endothelial cells, without interference from other tissues, as would occur in *in vivo* models. Human islets previously cultured with this platform have been shown to maintain their functionality^41^. In this three-dimensional assay, the device permits transit of media carrying immune cells. Therefore, T cell behavior can be directly visualized, allowing assessment and quantification of active T cell infiltration into single intact organoid structures. This is the first application of this vascularized platform to model antigen-specific autoimmune β-cell injury, extending prior work that focused on islet vascularization, function, and engraftment. For future studies, this setup enables readouts of T cell trafficking across endothelium, immune-synapse formation, cytokine flux, and endothelial activation^45^. These features cannot be captured using classic 2D/3D co-cultures.

Finally, infiltration was antigen-dependent and minimal with Ctrl Teffs in all organoids tested on the platform. PD-L1⁺ GINS formed fewer stable T cell contacts and displayed reduced T cell density, consistent with checkpoint-mediated attenuation of effector engagement (Fig. 4). These data indicate that the platform recapitulates physiologic trafficking and that antigen recognition is the primary driver of β-like organoid infiltration and injury.

In sum, GINS exhibit robust functional properties and reduced susceptibility to immune-mediated injury in comparison to same-donor iPSC-islets. In addition, GINS represent an engineerable β-like cell platform, and our findings demonstrate that checkpoint augmentation can further enhance their resistance to antigen-specific T cell-mediated cytotoxicity. Together, these results support the continued development of GINS as a platform for immune-protected cell replacement and advance their potential for both autologous and allogeneic applications without the need for extensive HLA editing. More broadly, this work highlights immunomodulatory engineering of β-like cells as a promising strategy to improve graft survival. However, an important limitation of this study is that these findings are derived from a single matched donor. While this design enables controlled comparisons by minimizing inter-donor variability, future studies across multiple donors will be essential to assess the reproducibility and generalizability of these observations.

## Materials and Methods

### Human samples

Human Gastric Stem Cells (hGSCs) and splenocytes were isolated from stomach^7^ and spleen (https://npod.org/wp-content/uploads/2021/05/OPPC-SOP-60.2-Isolation-of-Cells-from-Spleen-Thymus-and-Lymph-Nodes.pdf) of deceased donor obtained from Corpus Christi Medical Center, TX. Informed consent was obtained from the participants and or parents/guardian for these studies. Subject details are described in Supplementary Table 1.

### iPSC generation and differentiation

Splenocytes were reprogrammed to iPSC by the Integrated Genomics Operation (IGO) Core at the Memorial Sloan Kettering Cancer Center according to the following: The splenocytes-derived iPSC lines were generated based on an established PBMC reprogramming protocol. Briefly, splenocytes were cultured in StemSpan medium with erythroid expansion Supplement (Stem Cell Technology; #09655 and # 02692) for 7 days, and then 4 Sendai viral vector kit (ThermoFisher Scientific; A16517) expressing Oct3/4, Sox2, Klf4, or c-Myc were transduced to 2.5×10^5^ cells following the kit provided protocol. At day 20-25, 12 colonies were picked and expanded individually into 1 well of Matrigel coated 24-well plate in StemFlex culture medium (ThermoFisher Scientific, A3349401). During expansion, 3 clones were selected, and continue in culture for 8-10 passages to ensure stability of the lines. The iPSC clones were characterized using flow cytometry to assess a set of pluripotency markers including OCT4, SOX2 and NANOG (Fig. S1B). iPSCs were cultured on Matrigel and grown at 37°C with 5% CO2 and 20% O2 using StemFlex Basal Medium (Gibco A33493-01) with StemFlex supplement (Gibco A33492-01), provided by the Human Therapeutic Organoid Core at Weill Cornell Medicine. Cells were split every 3-4 days once reaching 80–90% confluence at a ratio of 1:12 using TrypLE dissociation reagent (Gibco) and replated with 10 μM Rock-inhibitor Y27632 dihydrochloride (Tocris). Differentiation to iPSC-derived β-like cells (iPSC-islets) was carried out using suspension-based protocol as previously described^18,19^, as the following:

- Embryoid body generation: Cells were incubated with the accutase solution for 8 min at 37°C, and cells were transferred to a 50 mL falcon tube washed twice using 40ml of DMEM-F12. 3 million cells were resuspended in 5 mL of hESC medium with 1 μM of ROCK inhibitor and plate them on one well of 6-well plate of ultra-low attachment cell culture plate. The plate was placed on an orbital shaker set up at 100rpm inside of incubator with 5% CO2 and 37°C. Overnight cells form embryoid bodies, which were fed for two days using hESC medium and ROCK inhibitor. After two days, the medium was removed, and it was replaced hESC medium without ROCK inhibitor. Upon 72 hrs, hESC medium was replaced with pancreatic differentiation medium for day zero. During differentiation protocol until day 14, medium was replaced every 24 h with fresh medium made during the day. After day 14 cells were fed every day after with fresh medium.

- Pancreatic differentiation: Day zero medium contains RPMI supplemented with 0.3 mM Chir99021, 100 μg/mL Activin A and 0.2% fetal bovine serum (FBS). For day 1 medium contains RPMI with 100 μg/mL Activin A, 0.03 mM Chir99021, 10 μg/mL bFGF and 0.2% FBS. For day 2 and 3 the backbone medium was SFD with 100 μg/mL Activin A. From day 4-6 the backbone medium was changed to DMEM-F12 with 50 ng/mL FGF7. Then medium was replaced from day 7-8 with DMEM high glucose (5 g/L) supplemented with 1:100 B27 without RA, 1X Glutamax, 0.25 mM ascorbic acid, 1:200 ITS-X, 50 ng/mL FGF7, 0.5 μM SANT-1, 1 μM Retinoic Acid, 100 nM LDN-193189 and 500 nM Phorbol. Medium for day 9-11 consist of DMEM high glucose (5 g/L) supplemented with 1:100 B27 without RA, 1X Glutamax, 0.25 mM ascorbic acid, 1:200 ITS-X, 2 ng/mL FGF7, 0.5 μM SANT-1, 0.1 μM Retinoic Acid, 200 nM LDN-193189 and 250 nM Phorbol. From day 12-14 the backbone medium was changed to MCDB131 supplemented with 20 mM glucose, 2% FBS, 1X Glutamax, 1:200 ITS-X, 10 μg/mL Heparin, 10 μM Zinc sulfate, 0.5 μM SANT-1, 0.05 μM Retinoic Acid, 200 nM LDN-193189, 1 μM T3 and 10 μM ALK5i II. From day 15-28 cells were fed every day after with medium that contains MCDB131 with 20 mM glucose, 2% FBS, 1X Glutamax, 1:200 ITS-X, 10 μg/mL Heparin, 10 μM Zinc sulfate, 200 nM LDN-193189, 1 μM T3, 10 μM ALK5i II and 100 nM GSIS XX. From day 29-40 cells were fed every day after with medium that contains MCDB131 with 20 mM glucose, 2% FBS, 1X Glutamax, 1:200 ITS-X, 10 μg/mL Heparin, 10 μM Zinc sulfate, 1 μM T3, 10 μM ALK5i II, 1 mM N-acetyl cysteine, 10 μM Trolox and 2 μM R428.

### Lentiviral packaging and titration

Lentiviral packaging and titration were carried out as previously described7.

### Human Gastric Stem Cells (hGSCs) culture and engineering with lentiviral infection for simultaneous expression of *MAFA*, *PDX1* and *PD-L1 (CD274)*

Human GSCs were cultured in hGSC medium as previously formulated^7^. To establish dual inducible cell lines, we first engineered Ngn3/ER-hGSCs as previously described^6^. Engineered cells were passaged in one well of six-well plate 24 hrs before lentiviral transduction. Cells were washed with DPBS 1x and overlaid with hGSC medium containing 10 μg/mL polybrene and 25 μL of a lentivirus carrying a construct containing genes encoding for *MAFA-T2A-PDX1 and/or MAFA-T2A-PDX1-T2A-PD-L1* under the TetON system (referred as *MAFA-PDX1* and/or *MAFA-PDX1-CD274*), with each gene separated by a sequence encoding for the T2A peptide (viral titer: ∼10^8^ TU/mL). Spinfection was then performed as follows: the cell culture with lentivirus was spun at 1,000*g* for 30 min at 37°C and then incubated at 37°C in a 7.5% CO_2_ incubator for 48 hrs. The medium was changed to hGSC medium containing 10 μg/mL blasticidin according to the selection marker incorporated into the construct for 2 weeks.

### Generation of monoclonal lines expressing *MAFA, PDX1* and or *PD-L1 (CD274)*

Human GSCs lines generated above were dissociated and plated in hGSC medium on a 10 cm plate and let them grow to form colonies. Single colonies were then picked and expanded and finally passaged into a 48-well plate. Each colony was then tested by immunofluorescence for the presence of MAFA upon induction with Tamoxifen and Dox simultaneously. Colonies of both lines including at least 90% of cells expressing MAFA, were selected. Monoclonal lines expressing *MAFA-PDX1* (clone A1) and or *MAFA-PDX1-CD274* (clone A6) were selected for the study. Monoclonal lines were used in the scRNA-seq, detection of T1D classic autoantigens and allogeneic/autologous immune assays. All the other experiments described include polyclonal lines of *MAFA-PDX1 and/or MAFA-PDX1-CD274*.

### GINS differentiation

GINS differentiation was carried out as follows: *Ngn3ER*/*MAFA-PDX1*-hGSCs and *Ngn3ER/MAFA-PDX1-CD274*-hGSCs were seeded 5 days before differentiation in 15 cm plate. To start differentiation, 1 μM 4-OH-TAM was added in hGSC medium and incubated for 2 days. The culture medium was then replaced with medium containing 50% hGSC medium and 50% serum-free basal medium^7^ with Dox 2 μg/mL to activate *MAFA-PDX1* and/or *MAFA-PDX1-CD274* expression. At day 4, GINS precursors were dissociated by 5-10 min TrypLE treatment and aggregated (typically 2.0-2.4 million cells per well) in AggreWell400 (STEMCELL Technologies, 34450) using the manufacturer’s recommended protocol. Medium was changed every 2-3 days. Aggregates were normally formed within 24 hrs.

### Static GSIS

GSIS for GINS organoids and iPSC-islets was carried out at day 16 and day 42 respectively. Ten to 20 organoids/line/biological replicate were sampled for each group. Organoids were washed with plain RPMI-1640 medium (RPMI, MyBio Source, MBS652918), and equilibrated in 3.3 mM glucose in RPMI for 2 hrs. Organoids were incubated in low-glucose RPMI for 1 h, and supernatant was collected. Organoids were then incubated in high-glucose (16.7 mM) RPMI for 1 h, and supernatant was collected. Secreted insulin was measured using the Stellux Chemi Human Insulin ELISA (ALPCO Diagnostics, 80-INSHU-CH10).

### Whole mount Immunofluorescence

GINS and mature iPSC-islets day 16 and 34 respectively were fixed in 4% paraformaldehyde for 15 min, washed in PBS + 0.2% Triton X-100 (PBS-T) 3×5 min, incubated with blocking buffer (10% Donkey serum in PBS-T) and stained with primary antibodies at the proper concentration overnight at 4°C. Samples were then washed in PBS + 0.1% Tween-20 (PBS-T20) and incubated in PBS-T20 with DAPI and respective secondary antibodies for 2 hrs at room temperature (r. t.). Samples were then washed in PBS-T and mounted in SlowFade Gold antifade reagent (Invitrogen S36936).

### Microscopy and imaging analysis

Images of immunostained organoids mounted in SlowFade Gold antifade reagent (see above) were taken with Leica TCS SP8 with 20 × NA 0.75 water objective, at the Microscopy and Image Analysis Core at Weill Cornell Medicine. The samples were excited at 405, 488, 552, and 638 nm for DAPI and Alexa Fluor 488, 546, and 647 respectively. Light was guided to the sample via an Acousto-Optical Beam Splitter. The emission light was guided via a size-adjustable pinhole, set at 115 µm and passed through Acousto-Optical Tunable Filters. Images were acquired as z-stacks with a step size of 1 µm. Images analysis was conducted using Fiji^46^. Wide-field images were taken with Nikon Eclipse Ti12 fluorescence microscope.

### Infiltration Index quantification

Quantification was performed using wide-field images. Fiji (National Institutes of Health) was used to measure fluorescent intensity of GFP^+^ T cells inside and outside the organoid in the following way (Fig. S3). First, the ROI was selected based on the organoid size. Threshold was adjusted on the organoid raw image using the software default algorithm. Next, the command “analyze particles” was used to select the ROI, similarly to what was previously done^47^. Parameters were adjusted to eliminate unwanted objects. Same fixed parameters were used for all the images. Next, background was subtracted from the T cell channel. And the ROI previously selected was applied to the image and duplicated to select the same ROI outside and adjacent to the organoid. Because the area outside the organoid is wider than the organoid area, four outside ROI were considered for each organoid at 12, 3, 6, 9 o’ clock (Fig. S3). Finally, integrated density (MGV*area of ROI) was extracted and used as fluorescent intensity to measure the total amount of fluorescence per organoid inside and outside the organoid. Infiltration Index was calculated as the ratio of the means between the two, similarly to previous study^48^. 95% Confidence Interval (CI) was then used to assess significance.

### Flow cytometry and Fluorescent Activated Cell Sorting (FACS)

GINS at different time points were dissociated in Accutase (Innovative Cell Technologies AT-104) for 15-20 min. Mature iPSC-islets at different time points were dissociated in Trypsin for 10 min. Fetal Bovine Serum (FBS) was then added to stop the reaction and dissociated cells were filtered through 40 μm nylon strains. Cells were then washed with 1x PBS. Extracellular staining was conducted for 15 min at r.t. in FACS buffer with conjugated antibodies. For intracellular staining, cells were fixed in 1.6% paraformaldehyde for 30 min at 37°C, washed with 1x PBS and stored in FACS buffer. Cells were then incubated with primary antibodies in saponin buffer (1x in dH2O, Biolegend, # 421002) for 30 min at r.t. Cells were then washed once with saponin and incubated with secondary antibodies for additional 30 min at r.t. Cells were then resuspended in FACS buffer and ran through either one of these flow cytometers: BD Symphony A5, Fortessa, or Fortessa X20 (according to availability at the Flow Cytometry Core Facility) in FACS buffer. FACS sorting was conducted using either one of these sorters: Influx, Symphony S6 I, Symphony S6 II (according to availability at the Flow Cytometry Core Facility). Flow cytometry analysis and FACS sorting were conducted at the Flow Cytometry Core at Weill Cornell Medicine.

### Single-cell RNA sequencing (scRNA-seq) from differentiated iPSC-islets and GINS organoids

iPSC-islets at day 34 and GINS organoids at day 16 post induction were collected and dissociated with Accutase (Innovative Cell Technologies AT-104) at 37°C for 15 and 30 min respectively. Fetal Bovine Serum (FBS) was then added to stop the reaction. Digested cells were then filtered through 40 μm nylon strains and resuspended in washed in 1x PBS and then resuspended in FACS buffer. Cell sorting at the Flow Cytometry Core Facility (see above) was then conducted to purify DAPI^-^ live cells. Purified samples were washed in PBS with 0.04% BSA (Millipore Sigma, A1595), then processed by 10x Genomics single-cell droplet sample preparation workflow at the Genomics Core Facility at Weill Cornell Medicine as previously described^49^.

### scRNA-seq analysis

scRNAseq datasets from GINS and iPSC-islets were analysed as previously described^7^. Importantly, Insulin-high cells were categorized by using Seurat unsupervised clustering on all cells in both GINS and iPSC-islets samples combined, and selecting the clusters with the highest INS gene expression.

### Generation of Avatar T cells

Autologous and allogeneic naïve CD8^+^ T cells were isolated using the EasySep Human Naïve CD8^+^ T cells isolation kit (Stem Cell, 18000). Autologous T cells were obtained from splenocytes of a dead donor, while allogeneic T cells were obtained from peripheral blood mononuclear cells (PBMCs). Naïve CD8^+^ T cells where then centrifuged at 350*g* and resuspended in cRPMI at a concentration of 2.5 x 10^5^ cells/mL. T cells were then resuspended with cRPMI containing human T cells activator CD3/CD28 Dynabeads at the concentration of 6.25 µL Dynabeads/mL, rhlL-2 (TECIN Teceleukin Ro 23-6019) at 50 IU/mL, and rhlL-7 (NSC nos. 780247) at 5 ng/mL and then incubated at 37°C and 5% CO2 for 48 hrs. They were then transduced with a lentiviral vector encoding a PPI-reactive T cell receptor (TCR) linked via a T2A sequence to a GFP reporter, or with GFP alone as a control. Following transduction, cRPMI containing protamine sulfate at 16 µg/mL, rhlL-2 at 100 IU/mL, and rhlL-7 at 10 ng/mL was added to the culture. Spinfection was then performed as follows: the cell culture with lentivirus was spun at 1,000*g* for 30 min at 37°C and then incubated at 37°C in a 7.5% CO2 incubator for 48 hrs. Media was changed every 48 hrs for 5 days. CD3/CD28 Dynabeads were then removed, and FACS was performed to sort transduced cells. Allogeneic T cells were transduced with lentiviruses carrying constructs encoding The MART-1 (melanoma antigen-specific) TCR^50^ and or HLA-A2 CAR (kind gift of the Levis Lab at University of British Columbia) and sorted using a BD FACS Melody Cell Sorter at the University of Florida (BD Biosciences, Franklin Lakes, NJ, USA), while autologous T cells were sorted at the Flow Core facility at Weill Cornell Medicine using either BD Influx cell sorter or BD Symphony S6 (BD Biosciences, Franklin Lakes, NJ, USA). cRPMI medium was prepared as follows: RPMI1640 without L-Glutamine (Corning 15-040-CV), 10% FBS (Genesee Scientific 25-550), 100x Penicillin/Streptomycin solution (Corning 30-002-Cl), 100x MEM Non-Essential Amino Acids (Gibco 11140-050), 1M HEPES Buffer (Gibco 15630-080), 100x Glutamax (Gibco 35050-061), 2-Mercaptoethanol (2 µL/500 mL Sigma-Aldrich M6250), 1M Sodium Hydroxide (Sigma-Aldrich S2770-100mL), 100 mM Sodium Pyruvate (Corning 25-000-Cl). Before each immune assay, avatar T cells were activated with 6.25 µL Dynabeads/mL, rhlL-2 at 50 IU/mL, and rhlL-7 at 5 ng/mL.

### Chromium release assay

Susceptibility of GINS and iPSC-islet lines to cell-mediated lysis by allogeneic T cell avatars was assessed using chromium-release assays in the format previously described by Chen *et al*^29,36^. GINS and iPSC-islets were stimulated with IFN-γ at 6 ng/mL for 24 hrs. Organoids were then dissociated as described above and when needed preproinsulin peptide PPI15-24 (ALWGPDPAAA)^14^ loading was conducted at 5 µM at 4°C for 30 min in 1x PBS. Cells were then radiolabelled with ^51^CrNa2O4 (Revvity, Waltham, MA, USA) at an activity of 1.48 x 10^5^ Bq/well for 4 hrs before washing twice with fresh cDMEM, prepared as followed: Dulbecco’s Modification of Eagle’s Medium (DMEM with 4.5 g/L glucose, L-glutamine and sodium pyruvate 1x, Corning 10-013-CV), 7.5% Bovine Serum Albumin (BSA, Corning A8412-100ML), 10% FBS (Genesee Scientific 25-550), 1M HEPES (Gibco 15630-080), 100x MEM Non-Essential Amino Acids (Gibco 11140-050), 100x Penicillin/Streptomycin solution (Corning 30-002-Cl). GINS and iPSC-islets were co-cultured with previously activated MART-1, PPI-reactive (1E6), and HLA-A2 CAR T cell avatars at 1:1, 5:1, and 10:1 effector:target (E:T) ratios for 48 hrs.

Following co-culture, the supernatants were removed, and along with 200 µL of 1x PBS used to wash the wells, were transferred into 6×50 mm lime glass tubes. The lysates of adherent cells were collected using a 2% SDS wash and transferred into separate tubes. ^51^Cr activity, measured in counts per minute, was assessed for both fractions on a Wizard 1470 automatic gamma counter (Revvity). The specific lysis of sBC was calculated as follows:

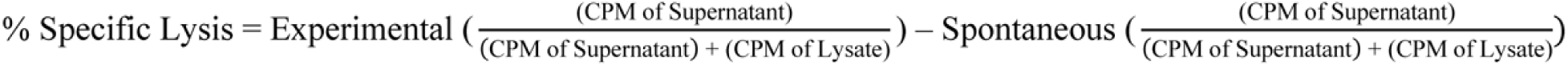

### Neutralization antibody experiment

Avatar T cells obtained from peripheral blood mononucleated cells, previously activated with rhlL-2 and rhlL-7 for 48 hrs, were incubated with PD-1 antibody for 20 min in FACS buffer at r.t., upon which they were added to the coculture with organoids previously primed with IFN-γ at 6 ng/mL for 24 hrs. Coculture was carried out for 48 hrs and cell death was measure through Chromium release assay, as specified above.

### Isolation and culture of Human Umbilical Vein Endothelial Cells (HUVECs)

Isolation of HUVECs and set up of the endothelial platform were carried out as previously described^41^ and as following: Human umbilical vein ECs (HUVECs) were isolated from discarded human umbilical cords, which was approved by the Weill Cornell Medicine Institutional Review Board (IRB). As previously described^51^ EC isolation was performed by digesting the umbilical cord vein using a collagenase-based solution. The ECs were then cultured with human EC culture medium, which is composed of M199, 20% fetal bovine serum, 2 mM glutamine, 15 mM Hepes, heparin (5 mg/mL), basic fibroblast growth factor (bFGF, FGF2) (10 ng/mL) (78134.1; STEMCELL Technologies), epidermal growth factor (EGF) (10 ng/mL) (78136; STEMCELL Technologies), insulin-like growth factor (IGF) (10 ng/mL) (78142.1; STEMCELL Technologies), and antibiotics.

### Immunochallenge of GINS and iPSC-islets with autologous T cells within the microfluidic device

GINS and iPSC-islets were stimulated with IFN-γ at 6 ng/mL for 24 hrs. Organoids were then stained with Vybrant™ DiI Cell-Labeling Solution for 30 min at 37°C, then co-mingled and loaded together with 260,000 of HUVECs and matrix in the microchambers, with a density of 50 organoids (2000 cells/organoid) per lane. To set up perfusion devices, HUVECs and organoids were resuspended in X-VIVO 10 serum-free hematopoietic stem cell medium (04-380Q; Lonza), mixed with bovine fibrinogen (341573-1GM; Sigma-Aldrich) to the final concentration of 2.5 mg/mL. The gelation of the cell/matrix mixture was induced by adding bovine thrombin to a final concentration of 1 U/mL. The microchambers used were the μ-Slide VI 0.4 (80606; ibidi) slides. Each chamber was loaded with 30 μL of EC-organoids/fibrin mixture. After complete gelation of the cell/fibrin mixture, two 3-mL syringes were attached to the ports on both sides of each lane, adding 2 mL of HUVEC tube-forming medium (described below) to one side (inlet) and 100 μL of tube-forming medium to the other side (outlet). The force of gravity to induce fluid transport from one side to the other was necessary for tube formation.

Vascularization was obtained within 48 hrs, upon which we added 200,000 T cells (either PPI Avatar autologous Teffs or Ctrl autologous Avatar Teffs) previously activated with rhlL-2 and rhlL-7 as described above. Incubation was carried out for 48 hrs.

Following, endothelial matrix including organoids and Avatar Teffs were dissociated with Nattokinase (Thermo Fisher 50-220-1212) and Accutase for 10-15 min and cells in suspension were recovered and filtered through 40 μm nylon strains. Cells were then stained live for anti-CD31, DAPI and Annexin V, as per manual instructions, and readout was performed by flow cytometry as previously described. Quantification was obtained through the normalization of the specific population taken into consideration (Annexin V^-^/DAPI^-^) to the sample incubated with Ctrl Teffs.

## Reagent list

Full list of antibodies used in this study is provided in Table 2.

**Table 1.** Donor information.

|  |  |
| --- | --- |
| CLASS I | A 2 A 3; B 7 B 44; BW4 Positive BW6 Positive; C 05 C 07 |
| CLASS II | DR 4 DR 15; DR51 51 DR52 N-Negative DR53 53; DQB1 6 DQB1 7; DQA1 01 DQA1 03; DPB1 04:01 DPB1 04:01; DPA1 01 DPA1 01 |

**Table 2.**
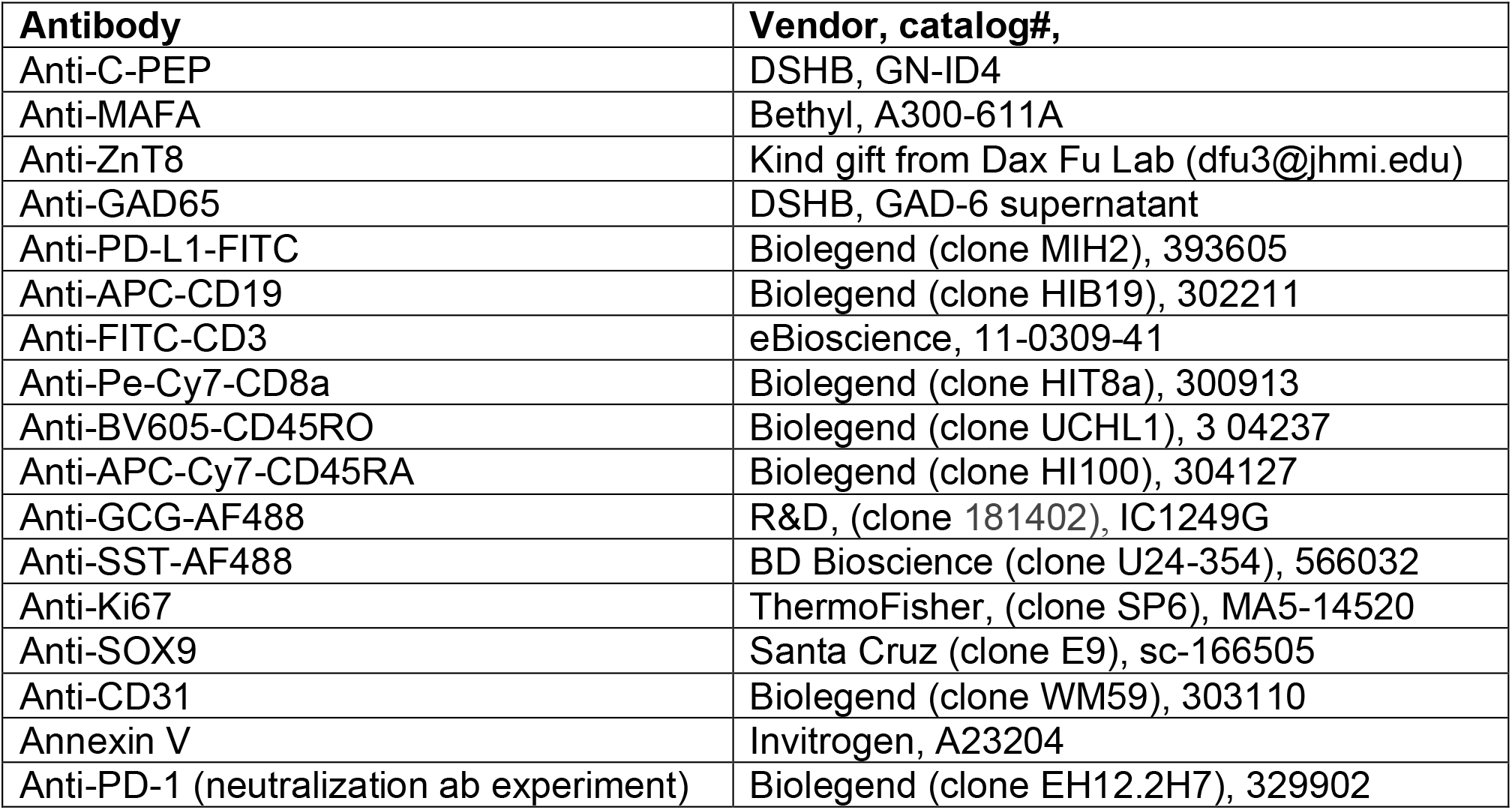
List of antibodies used.

## Acknowledgments

This research was supported by the National Institute of Health under the award number: R01DK106253, R01DK133701, R01DK133332, R01DK106191. The authors acknowledge the technical assistance of the Genomic Resources Core Facility at Weill Cornell Medicine and the Integrated Genomics Operation Core at the Memorial Sloan Kettering Cancer Center. The authors also acknowledge the facilities of the Flow Cytometry Core and the Microscopy and Image Analysis Core at Weill Cornell Medicine.

## Author contributions

Conceptualization: AAD, QZ, XH. Characterization of iPSC-islets, GINS and PD-L1^+^GINS: AAD, YL, VP, MZ. Genomic analysis: AAD, RN, XH. Chromium release assay for allogeneic immune assay: MEB and AAD. Autologous immune assay with the endothelial platform: AAD, ZF, BP, YK. Data discussion and interpretation: AAD, XH, RJC, TB, SR. Manuscript original draft: AAD and XH.

## Supplementary material

**Fig. S1.**
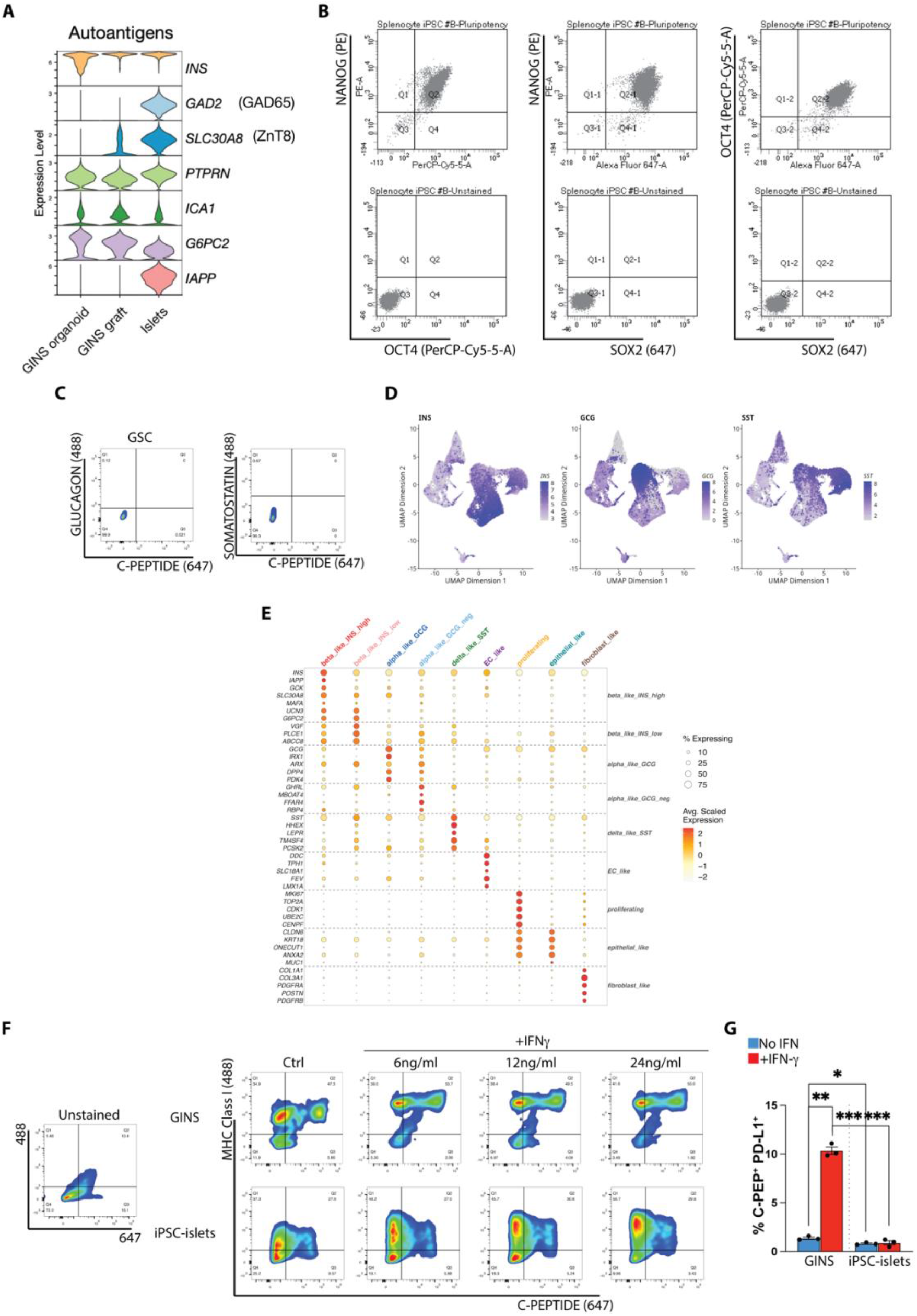
Supporting Figure for Fig. 1 and 2. (A) Violin plots from scRNA-seq of classic T1D autoantigens obtained either from GINS organoids, GINS graft or human islets. (B) Flow cytometry of stem cell markers NANOG, OCT4, SOX2 expression upon reprogramming of splenocytes into iPSCs. Bottom row shows unstained samples. (C) Flow cytometry of GSC for C-PEP, GCG, SST (negative control for Fig. 1E). (D) UMAP of GINS organoids and or iPSC-islets showing Insulin (INS), Glucagon (GCG), Somatostatin (SST) expression. (E) Expression of selective markers for each specific cell type identified. (F) Flow cytometry of MHC class I and C-PEP expression upon incubation with different doses of IFN-γ for 24 hrs. (G) Flow cytometry quantification of PD-L1^+^ C-PEP^+^ fraction at baseline and upon 24 hrs IFN-γ (6 ng/mL) treatment. All graphs depict mean and SEM. All quantifications are based on n=3 independent differentiations (unless differently specified) and were analyzed by paired t test. ns, not significant; *p < 0.05; **p < 0.01; ***p < 0.001; ****p < 0.0001.

**Fig. S2.**
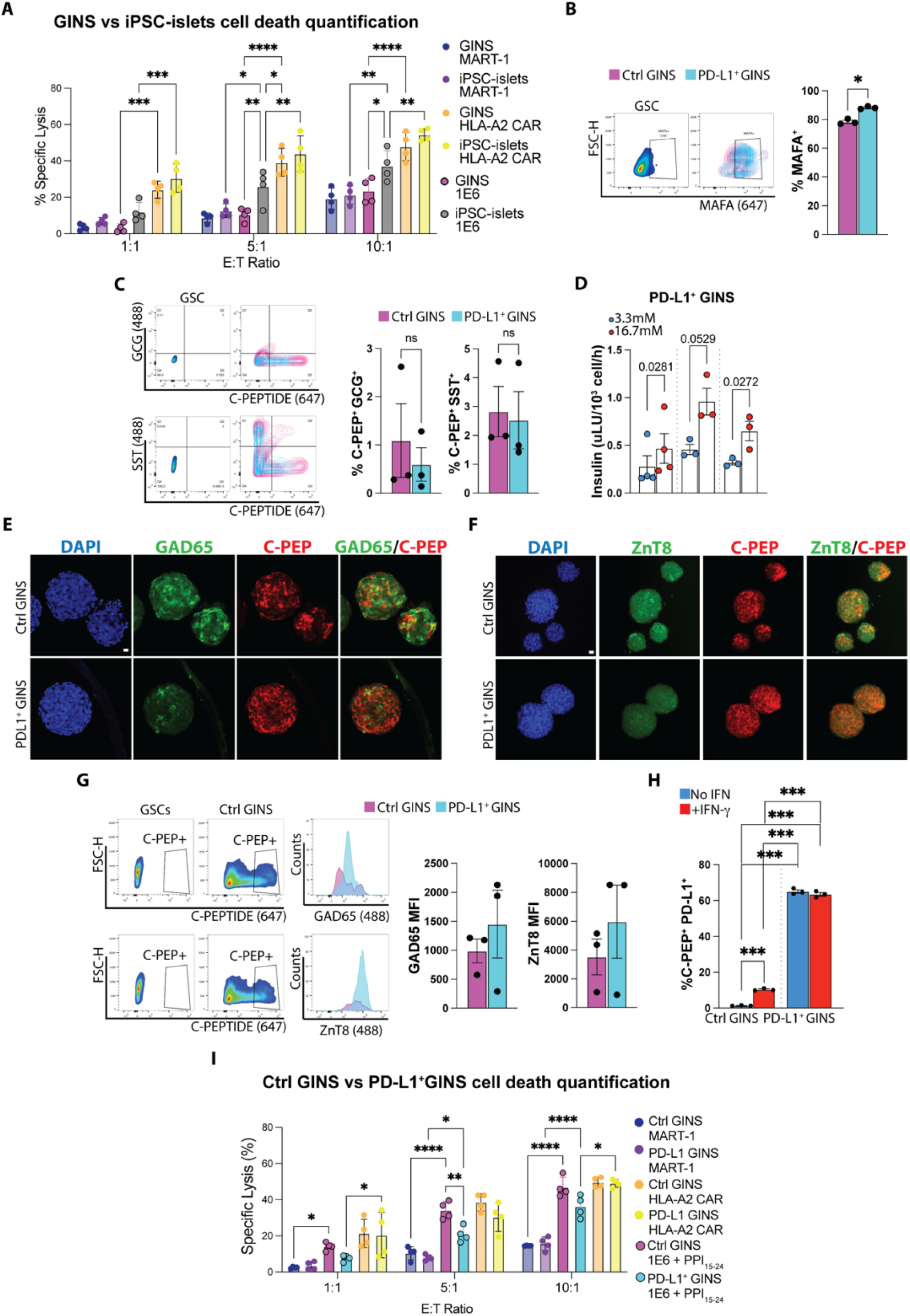
Supporting Figure for Fig. 2-3. (A) Cell death quantification measured as % specific lysis of GINS (Day 23) and iPSC-islets (Day 43) upon incubation with allogeneic T cells specific for preproinsulin peptide (PPI-Avatar clone 1E6) including negative and positive controls. (B) Flow cytometry plots and relative quantification of MAFA expression in Ctrl GINS and PD-L1^+^ GINS (*n* = 3 biological replicates, Days 21-28). (C) Flow cytometry quantification of Glucagon (GCG) and Somatostatin (SST) in combination with C-PEP expression on day 23 for GINS and PD-L1^+^ GINS (*n* = 3 biological replicates, Days 21-28). (D) GSIS of PD-L1^+^ GINS at day 16. (E) Immunofluorescence staining for DAPI (blue), C-PEP (red) and GAD65 (green) at day 19 for GINS and PD-L1^+^ GINS. Scale bar represents 10 μm. (F) Immunofluorescence staining for DAPI (blue), C-PEP (red) and ZnT8 (green) at day 19 for GINS and PD-L1^+^ GINS. Scale bar represents 20 μm. (G) Flow cytometry plots and quantification of C-PEP, GAD65 and ZnT8 expression. (H) Flow cytometry quantification of C-PEP^+^ PD-L1^+^ fraction at baseline and upon 24 hrs IFN-γ (6 ng/mL) treatment. (I) Cell death quantification measured as % specific lysis of GINS (Day 23) and iPSC-islets (Day 43) upon incubation with allogeneic T cells specific for preproinsulin peptide (PPI-Avatar clone 1E6) including negative and positive controls. All graphs show mean and Standard Error of the Mean (SEM). All quantifications are based on n=3 independent differentiations (unless differently specified) and were analyzed by paired t test. ns, not significant; *p < 0.05; **p < 0.01; ***p < 0.001; ****p < 0.0001.

**Fig. S3.**
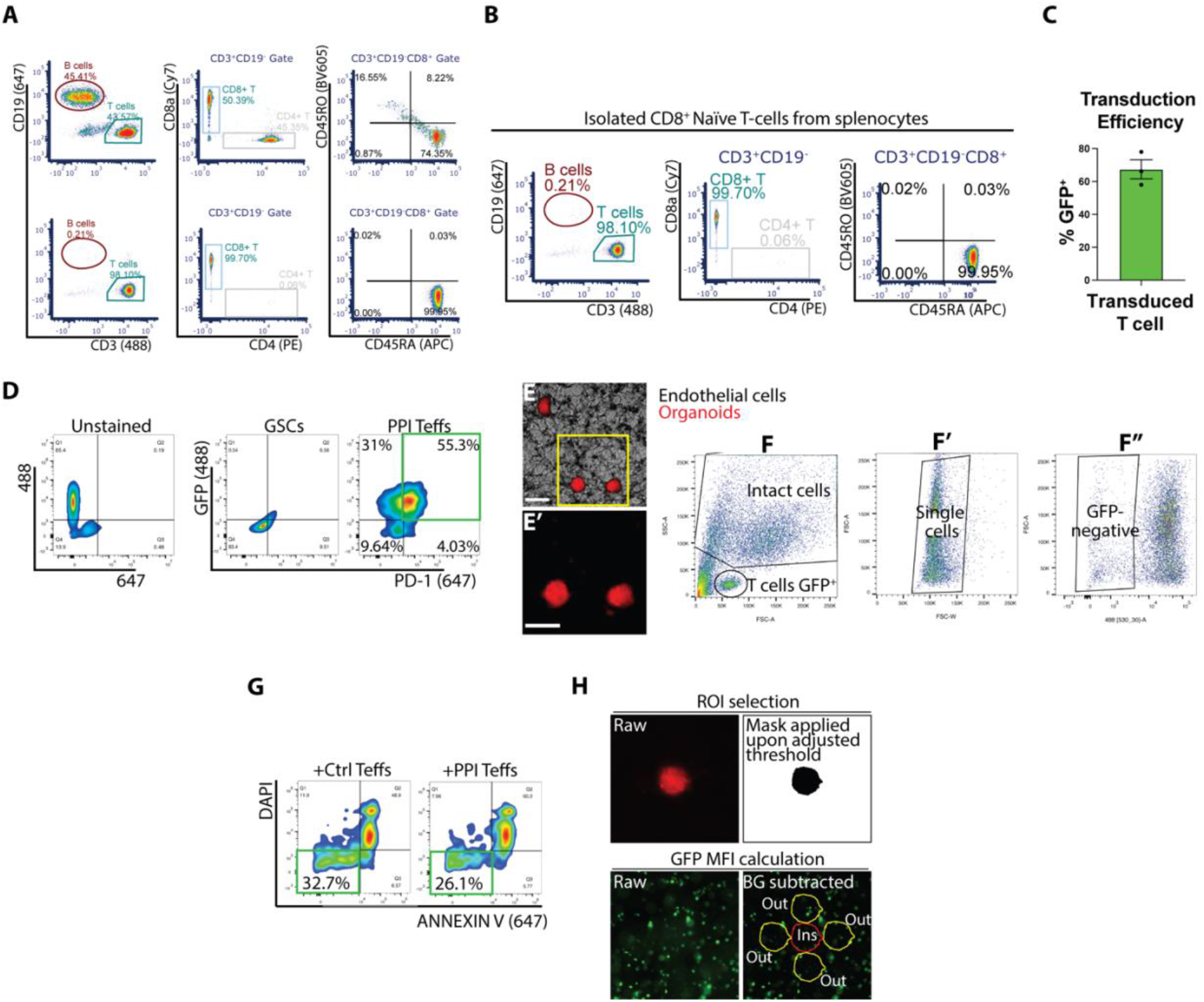
Supporting Figure for Fig. 4. (A) Flow cytometry of CD19, CD3, CD4, CD8a, CD45RO, CD45RA expression before isolation of autologous CD8^+^ naïve T cells. (B) Transduction efficiency (measured as GFP^+^) cells of the construct carrying the PPI TCR (1E6) linked to GFP through the T2A peptide (n=3 independent viral transduction). (C) Flow cytometry of PD-1 expression upon Teffs incubation with rhlL-2, rhlL-7 and CD28/CD3 beads. (D-D’) Vascularized iPSC-islet organoids on the endothelial microfluidic device, scale bars represent 100 μm. (E-E’) Representative flow cytometry plots illustrating the gating strategy to analyze cell death in the samples. (F) Flow cytometry of iPSC-islet cell viability upon T cell incubation measured using DAPI and Annexin V. (G) Example image used for quantification (see Material & methods); BG=Background. All graphs show mean and SEM. All quantifications are based on n=3 independent differentiations (unless otherwise specified) and were analyzed by paired t test. ns, not significant; *p < 0.05; **p < 0.01; ***p < 0.001; ****p < 0.0001.

## Bibliography

1. Vecchio, I., Tornali, C., Bragazzi, N. L. & Martini, M. The discovery of insulin: An important milestone in the history of medicine. Frontiers in Endocrinology vol. 9 (2018).

2. Efrat, S. Cell replacement therapy for type 1 diabetes. Trends Mol. Med. 8, 334–340 (2002).

3. Bellin, M. D. et al. Potent induction immunotherapy promotes long-term insulin independence after islet transplantation in type 1 diabetes. Am. J. Transplant. 12, 1576–1583 (2012).

4. Ruiz, R. & Kirk, A. D. Long-Term ToxiciTy of immunosuppressive Therapy.

5. Reichman, T. W. et al. Stem Cell-Derived, Fully Differentiated Islets for Type 1 Diabetes. N. Engl. J. Med. 393, 858–868 (2025).

6. Dattoli, A. A., Kelemen, Y. & Huang, X. Reprogramming of different cell lineages into functional B-cell substitutes. Cell. Reprogram. (2025).

7. Huang, X. et al. Stomach-derived human insulin-secreting organoids restore glucose homeostasis. Nat. Cell Biol. 25, 778–786 (2023).

8. Xu, L. et al. Expression of a mutant CD47 protects against phagocytosis without inducing cell death or inhibiting angiogenesis. Cell Reports Med. 5, (2024).

9. Carlsson, P.-O. et al. Survival of Transplanted Allogeneic Beta Cells with No Immunosuppression. N. Engl. J. Med. (2025) doi:10.1056/nejmoa2503822.

10. Gravina, A. et al. Synthetic immune checkpoint engagers protect HLA-deficient iPSCs and derivatives from innate immune cell cytotoxicity. Cell Stem Cell 30, 1538–1548.e4 (2023).

11. Sun, C., Mezzadra, R. & Schumacher, T. N. Regulation and Function of the PD-L1 Checkpoint. Immunity 48, 434–452 (2018).

12. Han, Y., Liu, D. & Li, L. PD-1/PD-L1 pathway: current researches in cancer. Am. J. Cancer Res. 10, 727–742 (2020).

13. Santini-González, J. et al. Human stem cell derived beta-like cells engineered to present PD-L1 improve transplant survival in NOD mice carrying human HLA class I. Front. Endocrinol. (Lausanne). 13, 1–14 (2022).

14. Castro-Gutierrez, R., Alkanani, A., Mathews, C. E., Michels, A. & Russ, H. A. Protecting Stem Cell Derived Pancreatic Beta-Like Cells From Diabetogenic T Cell Recognition. Front. Endocrinol. (Lausanne). 12, 1–12 (2021).

15. Arvan, P., Pietropaolo, M., Ostrov, D. & Rhodes, C. J. Islet autoantigens: Structure, function, localization, and regulation. Cold Spring Harb. Perspect. Med. 2, 1–20 (2012).

16. Williams, C. L. & Long, A. E. What has zinc transporter 8 autoimmunity taught us about type 1 diabetes? Diabetologia 62, 1969–1976 (2019).

17. Trabucchi, A. et al. Development of an immunoassay for the simultaneous detection of GADA and ZnT8A in autoimmune diabetes using a ZnT8/GAD65 chimeric molecule. Front. Immunol. 14, (2023).

18. Leavens, K. F. et al. Generation of a fluorescent mNeonGreen insulin reporter line in the H1 (WA01) hESC background. Stem Cell Res. 81, 103559 (2024).

19. Cardenas-Diaz, F. L. et al. Modeling Monogenic Diabetes using Human ESCs Reveals Developmental and Metabolic Deficiencies Caused by Mutations in HNF1A. Cell Stem Cell 25, 273–289.e5 (2019).

20. Richardson, S. J. et al. Islet cell hyperexpression of HLA class I antigens: a defining feature in type 1 diabetes. Diabetologia 59, 2448–2458 (2016).

21. Demine, S. et al. Pro-inflammatory cytokines induce cell death, inflammatory responses, and endoplasmic reticulum stress in human iPSC-derived beta cells. Stem Cell Res. Ther. 11, 1–15 (2020).

22. Moon, J. W. et al. IFNγ induces PD-L1 overexpression by JAK2/STAT1/IRF-1 signaling in EBV-positive gastric carcinoma. Sci. Rep. 7, 1–13 (2017).

23. Newby, B. N. et al. Type 1 interferons potentiate human cd8+ t-cell cytotoxicity through a stat4-and granzyme b-dependent pathway. Diabetes 66, 3061–3071 (2017).

24. Robbins, P. F. et al. Single and Dual Amino Acid Substitutions in TCR CDRs Can Enhance Antigen-Specific T Cell Functions. J. Immunol. 180, 6116–6131 (2008).

25. Freed-Pastor, W. A. et al. The CD155/TIGIT axis promotes and maintains immune evasion in neoantigen-expressing pancreatic cancer. Cancer Cell 39, 1342–1360.e14 (2021).

26. Sordo-Bahamonde, C. et al. Beyond the anti-PD-1/PD-L1 era: promising role of the BTLA/HVEM axis as a future target for cancer immunotherapy. Mol. Cancer 22, 1–15 (2023).

27. Deuse, T. et al. Hypoimmunogenic derivatives of induced pluripotent stem cells evade immune rejection in fully immunocompetent allogeneic recipients. Nat. Biotechnol. 37, 252–258 (2019).

28. Pereira, B. I. et al. Senescent cells evade immune clearance via HLA-E-mediated NK and CD8+ T cell inhibition. Nat. Commun. 10, (2019).

29. Brown, M. E. et al. Human stem cell- - derived β cells expressing an optimized CD155 reduce cytotoxic immune cell function for application in type 1 diabetes. 1–17 (2025) doi:10.1126/sciadv.adx9755.

30. Medina-serpas, M. A. et al. Resource Spatial transcriptomics from pancreas and local draining lymph node tissue reveals a lymphotoxin- β signature in human type 1 diabetes ll Spatial transcriptomics from pancreas and local draining lymph node tissue reveals a lymphotoxin- β signatur. CellReports 45, 117144 (2026).

31. Beattie, J. H. et al. Metallothionein overexpression and resistance to toxic stress. Toxicol. Lett. 157, 69–78 (2005).

32. Rosa, A. C., Corsi, D., Cavi, N., Bruni, N. & Dosio, F. Superoxide Dismutase Administration: A Review of Proposed Human Uses. Molecules 26, (2021).

33. Cheng, Y.-Y. et al. Reactive astrocytes increase expression of proNGF in the mouse model of contused spinal cord injury. Neurosci. Res. (2019) doi:10.1016/j.neures.2019.07.007.

34. Chen, Q. et al. LTβR-RelB signaling in intestinal epithelial cells protects from chemotherapy-induced mucosal damage. Front. Immunol. **Volume** 15, (2024).

35. D’Oliveira Albanus, R., et al. Integrative single-cell multi-omics profiling of human pancreatic islets identifies T1D-associated genes and regulatory signals. Cell Rep. 44, (2025).

36. Chen, J., Grieshaber, S. & Mathews, C. E. Methods to assess beta cell death mediated by cytotoxic T lymphocytes. J. Vis. Exp. 1–4 (2011) doi:10.3791/2724.

37. Furuyama, K. et al. Diabetes relief in mice by glucose-sensing insulin-secreting human α-cells. Nature doi:10.1038/s41586-019-0942-8.

38. Li, Y. et al. Humanized Mice Reveal New Insights Into the Thymic Selection of Human Autoreactive CD8+ T Cells. Front. Immunol. 10, 1–16 (2019).

39. Greiner, D. L. et al. Humanized mice for the study of type 1 and type 2 diabetes. Ann. N. Y. Acad. Sci. 1245, 55–58 (2011).

40. Koning, M. et al. Vasculogenesis in kidney organoids upon transplantation. npj Regen. Med. 7, 1–16 (2022).

41. Li, G. et al. Vascularization of human islets by adaptable endothelium for durable and functional subcutaneous engraftment. Sci. Adv. 11, 1–13 (2025).

42. Lertkiatmongkol, P., Liao, D., Mei, H., Hu, Y. & Newman, P. J. Endothelial functions of platelet/endothelial cell adhesion molecule-1 (CD31). Curr. Opin. Hematol. 23, (2016).

43. Vermes, I., Haanen, C., Steffens-Nakken, H. & Reutellingsperger, C. A novel assay for apoptosis Flow cytometric detection of phosphatidylserine expression on early apoptotic cells using fluorescein labelled Annexin V. J. Immunol. Methods 184, 39–51 (1995).

44. Hu, X. et al. Human hypoimmune primary pancreatic islets avoid rejection and autoimmunity and alleviate diabetes in allogeneic humanized mice. Sci. Transl. Med. 15, 1–16 (2023).

45. Yengej, F. A. Y., Wal, M. M. Van Der & Anderson, G. T cell interaction with activated endothelial cells primes for tissue-residency. 1–13 (2022) doi:10.3389/fimmu.2022.827786.

46. Schindelin, J., et al. Fiji: an open-source platform for biological-image analysis. Nat. Methods 9, 676–682 (2012).

47. Dattoli, A. A. et al. Asymmetric assembly of centromeres epigenetically regulates stem cell fate. J. Cell Biol. 219, e201910084 (2020).

48. Dattoli, A. A. et al. Domain analysis of the Nematostella vectensis SNAIL ortholog reveals unique nucleolar localization that depends on the zinc-finger domains. Sci. Rep. 5, 1–20 (2015).

49. Gu, W. et al. SATB2 preserves colon stem cell identity and mediates ileum-colon conversion via enhancer remodeling. Cell Stem Cell 29, 101–115.e10 (2022).

50. Johnson, L. A. et al. Gene Transfer of Tumor-Reactive TCR Confers Both High Avidity and Tumor Reactivity to Nonreactive Peripheral Blood Mononuclear Cells and Tumor-Infiltrating Lymphocytes. J. Immunol. 177, 6548–6559 (2006).

51. Baudin, B., Bruneel, A., Bosselut, N. & Vaubourdolle, M. A protocol for isolation and culture of human umbilical vein endothelial cells. Nat. Protoc. 2, 481–485 (2007).

