## Supplementary figures and images for "Generation of hypoimmunogenic gastric insulin-secreting organoids"

### Fig. S1

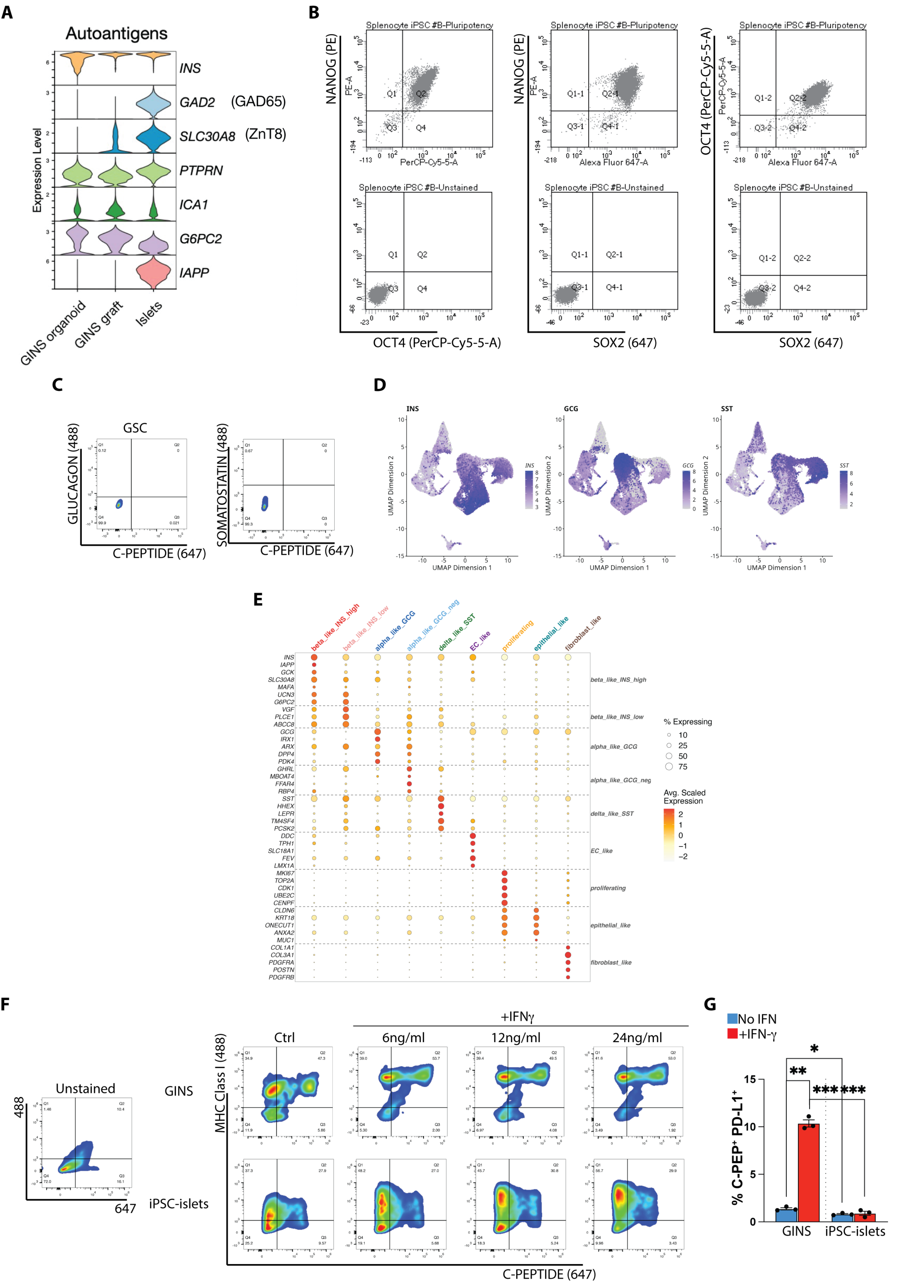

### Fig. S2

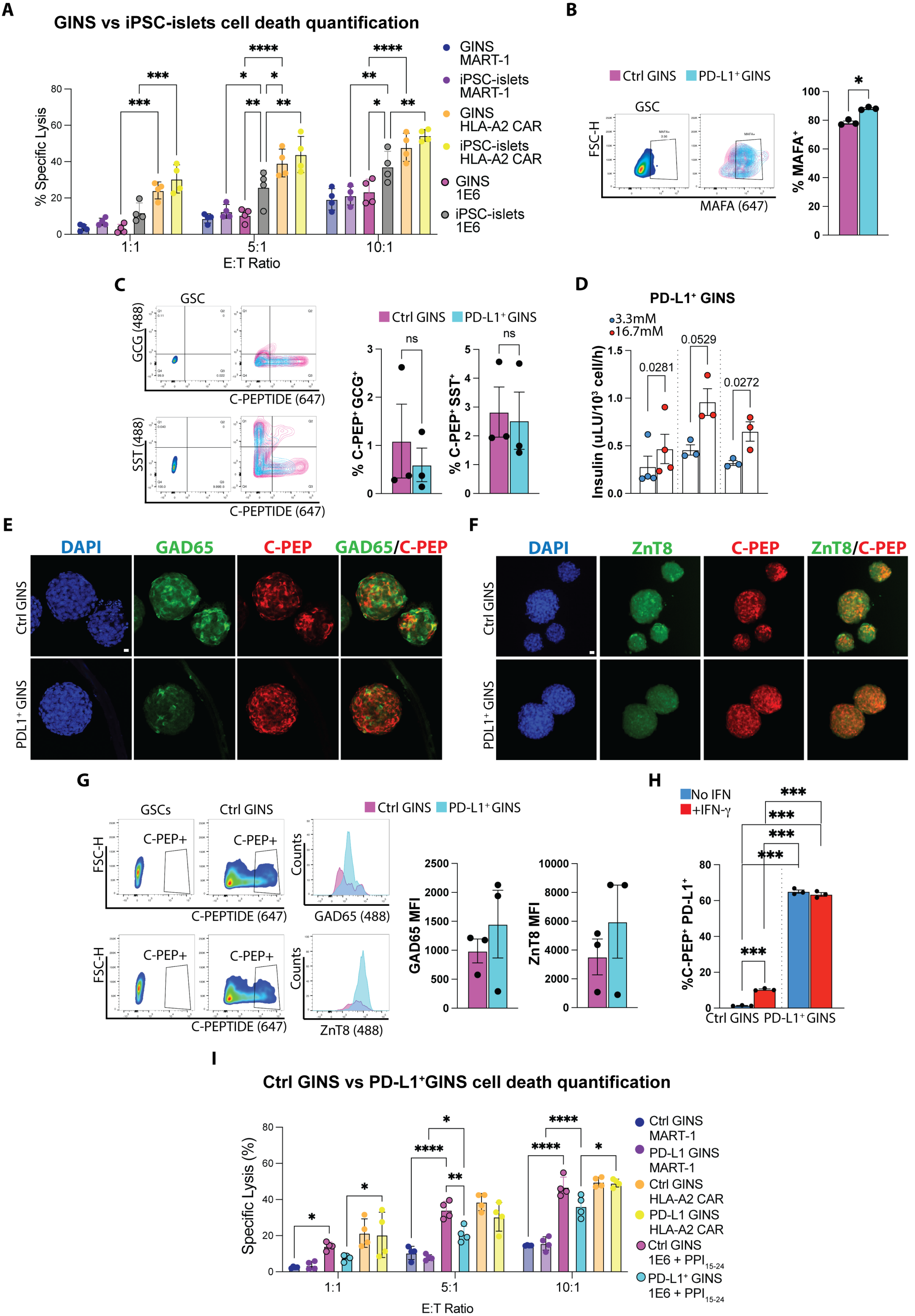

### Fig. S3

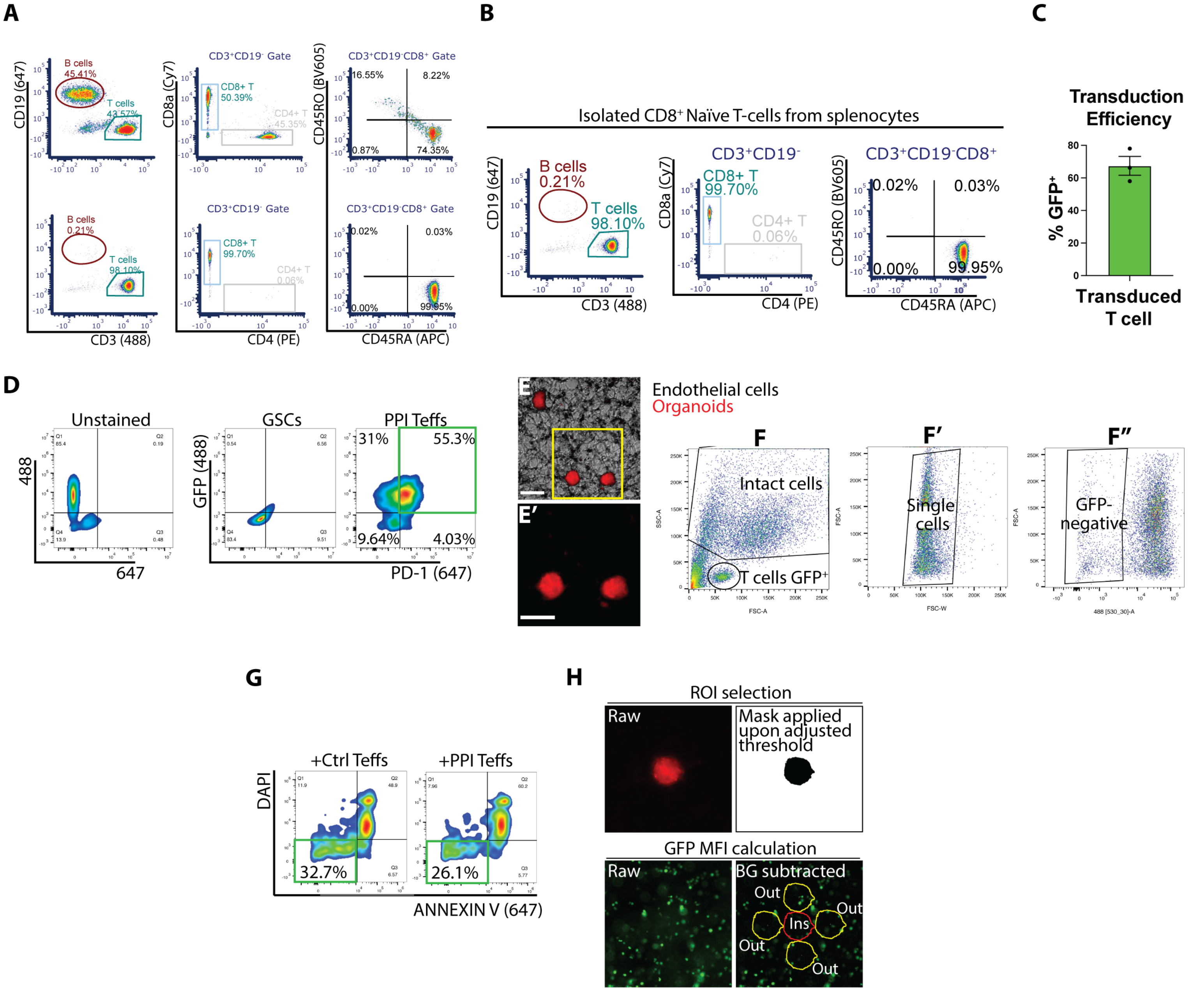
